# Intestinal Stem and Progenitor Cells Exhibit Distinct Adaptive Responses to Inflammatory Stress in IBD

**DOI:** 10.64898/2025.12.11.691866

**Authors:** Brinda Balasubramanian, Shivam Patel, Louis Gall, Nick Hannan, William Dalleywater, Joerg Huelsken, Carmen Pin, Gordon Moran, Paloma Ordóñez-Morán

## Abstract

**Background:** Intestinal epithelial stem cells (SCs) and their transit-amplifying (TA) progeny are critical for mucosal repair and regeneration. However, their behaviour under chronic inflammatory conditions, such as those observed in Inflammatory Bowel Disease (IBD), remains incompletely understood.

**Methods:** We investigated the impact of chronic inflammation on intestinal stem/progenitor cells by integrating bulk RNA sequencing from the largest IBD biopsy cohort to date with single-cell transcriptomic analysis and experimental assays using patient-derived intestinal organoids.

**Results:** Active inflammation was associated with a reduction in canonical LGR5⁺ intestinal stem cells and a concurrent expansion of OLFM4⁺ populations, consistent with an inflammation-induced epithelial repair program. Notably, SC/TA cells from both inflamed and non-inflamed IBD tissues exhibited persistent transcriptional changes that were distinct from those in healthy controls. Single-cell analysis identified transcriptionally heterogeneous SC/TA subpopulations, including a previously uncharacterized inflammation-associated cluster enriched in immune signalling pathways. Pseudotime trajectory analysis demonstrated a shift in differentiation toward deep crypt secretory (Paneth-like) cell lineages under inflammatory conditions.

**Conclusions:** Chronic intestinal inflammation reshapes the epithelial stem and progenitor cell compartment, promoting altered differentiation and the emergence of immune-responsive epithelial states. These findings highlight the plasticity of the human intestinal epithelium in IBD and point to new avenues for therapeutic strategies aimed at maintaining epithelial integrity during chronic inflammation.

## Introduction

Inflammatory Bowel Disease (IBD) is increasingly recognized by the World Health Organization as a major global health concern, especially given its rising prevalence in high-income countries and growing incidence among younger populations ^1–3^. IBD encompasses chronic, relapsing-remitting inflammatory disorders of the gastrointestinal tract, with Crohn’s disease (CD) and Ulcerative Colitis (UC) being the primary subtypes. These conditions are driven by an abnormal immune response directed at the intestinal lining, resulting in symptoms such as abdominal pain, diarrhea, fatigue, and weight loss. Over time, approximately 70% of individuals with CD and 20–30% of those with UC require surgical intervention due to complications like strictures, fistulas, abscesses, or failure of medical therapy and malignancy ^2,3^. Although the exact aetiology of IBD remains unresolved, current evidence supports a multifactorial origin involving genetic susceptibility, environmental triggers, and immune system dysregulation.

During chronic inflammation, epithelial cells are persistently exposed to pro-inflammatory cytokines and oxidative stress, leading to the activation of signalling pathways such as Hippo-YAP1 and Wnt/β-catenin, which are crucial for maintaining intestinal homeostasis^4–6^. This inflammatory environment compromises the gut barrier by altering epithelial function and increasing tissue permeability, thereby intensifying immune responses ^7,8^. These changes negatively impact the proliferation of intestinal stem cells (SC) and transit-amplifying progenitors (TA), while also promoting cell death, ultimately weakening epithelial integrity ^9^. SC, which reside at the intestinal crypts, are essential for sustaining epithelial renewal by generating TA that divide and differentiate into the diverse mature cell types lining the intestine ^10–12^. It is unclear how SC and TA (SC/TA) attempt to repair or adapt to the inflammatory environment, or what specific signals drive their response in a timely manner in healthy and IBD tissue. It is particularly relevant to know how these cells adapt under stress: whether they shift towards rapid proliferation to replace damaged cells, lean towards self-renewal to preserve the stem cell pool or undergo differentiation.

Bestrophin-4 (BEST4)⁺ cells, are a subpopulation of mature absorptive epithelial cells characterized by the expression of the ion channel BEST4^13^. These cells have an important physiological relevance because their unique gene expression profile shows that they have a specialized functions in pH regulation, electrolyte transport, antimicrobial defense, and immune communication^14^. BEST4^+^ cells are emerging as an important cell type in IBD and their loss in inflammation has been reported^14–16^, Disruptions among BEST4⁺ cells could impact barrier function, antimicrobial defense, and local immune environments, processes highly relevant to IBD pathophysiology. However, the underlying mechanisms, functional consequences, and potential for barrier permeability and regeneration remains unclear. One of the main reasons for limited progress in this area is the absence of BEST4⁺ cells in mice^17–20^. Integrating them into our human studies of SC/TA populations is relevant for better understanding the origins of BEST4^+^ as they may arise from distinct stem or transit-amplifying cells and understanding their role during chronic inflammation. Overall, the functional relevance of SC/TA and BEST4^+^ cells and its dynamic changes in IBD could reveal key insights into tissue repair, novel therapeutical approaches and the understanding of IBD progression.

In this study, we set out to identify the dynamic molecular response of intestinal SC, TA and BEST4^+^ epithelial cell types during inflammation by comparative analysis of omics data and experimental approaches using IBD patients tissue data, DSS-treated UC mouse models and IBD patient-derived organoids (PDOs). Using our multimodal approach, we assessed changes in cell type markers, identified biological processes, and pathway modulations associated with inflammation. Our approach offers the first evidence in a controlled setting to trace molecular signatures and adaptive responses of distinct epithelial molecular clusters in relation to inflammation severity, providing valuable insights into how inflammatory processes disrupt SC/TA epithelial function and contribute to IBD progression. Modulation of these cell types reflects a shift in stemness and differentiation pathways, where specific OLFM4⁺ SC/TA cells increasingly remain in a proliferative state under heightened inflammation, rather than progressing toward full epithelial maturation. Moreover, BEST4^+^ mature cells showed a slight increase in low-inflammation tissue, however, their numbers were significantly downregulated under high inflammatory conditions. These cells may play a regulatory role in maintaining epithelial homeostasis and their reduction during severe inflammation could contribute to disease progression in IBD. This epithelial remodelling highlights a potential pathogenic imbalance between stemness and differentiation in both inflamed and non-inflamed IBD epithelium. Furthermore, we detected that the differentiation trajectory of IBD stem cells is skewed toward the deep crypt secretory cell lineage, also referred to as colonic Paneth-like cells, indicating an aberrant differentiation process. Collectively, our findings demonstrate for the first time how intestinal stem cell and progenitor epithelial cell types actively engages in adaptive responses to inflammation, balancing proliferation and differentiation processes that are critical for tissue repair and epithelial.

## Results

### Profiling epithelial cell heterogeneity based on inflammation severity

To understand the epithelial cell heterogeneity under varying degrees of inflammation, we investigated the cell type distribution based on macroscopic appearance of IBD tissue. To that end, we utilized bulk-RNA sequencing (RNAseq) data from the largest cohort of IBD patients’ biopsies analysed to date (IBD Plexus Study of a Prospective Adult Research Cohort, SPARC IBD) (UC=1,097, CD=2,138) of the Crohn’s & Colitis Foundation IBD Plexus program which includes clinical, genetic, molecular, environmental, and patient-reported outcomes from individuals with CD or UC^21^ (Figure S1A).

We employed deconvolution analysis using CIBERSORTx to quantitatively estimate the epithelial, immune, and stromal cells type proportion with a signature matrix constructed using single cell-RNAseq (sc-RNAseq) data from human intestine as reference^15^ (Figure 1A and S1B). The overall quantitative relative abundance of cycling TA, and immature enterocytes (2) pool involved mainly in metabolic processes, decreased with increasing inflammation severity reflecting an altered capacity within the epithelial lining. However, there was an increase in immature enterocytes (1), which are involved in tight junction repair during inflammation^22^. This rise may be a compensatory mechanism, aiming to restore barrier integrity despite the diminished support from the TA pool. In parallel, M cells, key players in mucosal immune surveillance, were significantly enriched in inflamed tissues, consistent with heightened immune activity (Figure 1B and S1C). Well-characterized SC, TA, BEST4^+^ cell type markers and/or Wnt pathway target genes, such as *LGR5*, *AXIN2*, *EPHB2, ASCL2, PROM1*, *LRIG1* and *BEST4*, were significantly lower expressed in inflammation vs normal or possible inflammation of UC (Figure 1C, D) and CD (Figure S1D, E). In contrast, *OLFM4*, a key SC/TA marker strongly associated with cell proliferation and rapid cellular renewal ^23^, increased as inflammation intensified (Figure 1C, D and S1D, E). Clusterin (*CLU*) ‘revival stem cell’ pool marker involved in tissue regeneration after injury^24^ increased with inflammation severity (Figure 1C and S1D). This suggested an adaptive response, where OLFM4^+^ proliferative cells in IBD tissue aim to counterbalance the loss of epithelial integrity caused by inflammation.

**Figure 1.**
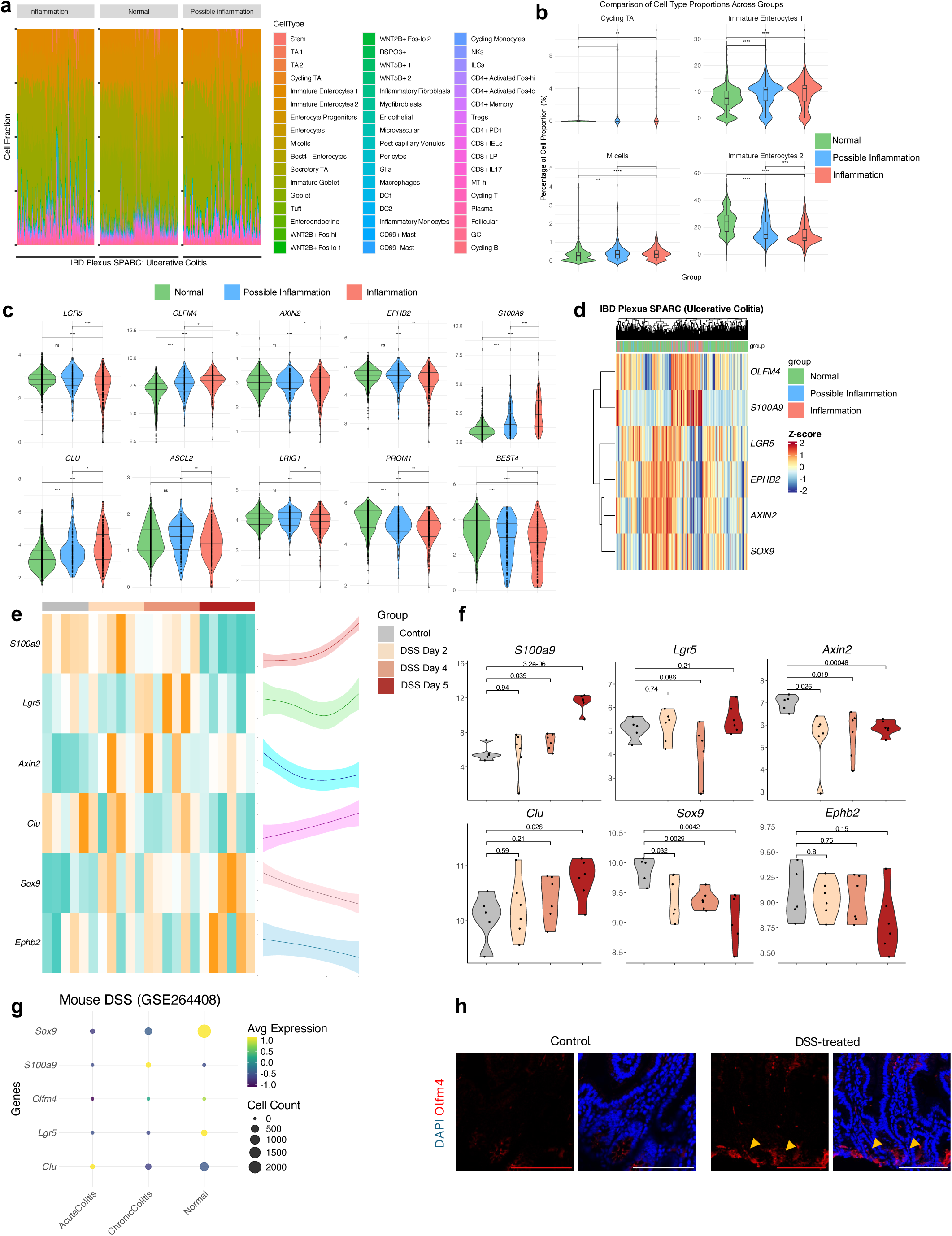
Epithelial cell heterogeneity profile varies with inflammation severity in IBD. (A) Bar plots showing the relative proportions of epithelial cell types stratified by endoscopic inflammation scores in SPARC IBD-UC samples. (B) Violin plots depicting significant differences in cell type proportions across macroscopic appearance groups (normal, possible inflammation, inflammation). (C) Violin plots of representative epithelial gene markers across the same macroscopic groups. (D) Heatmap showing unsupervised clustering of epithelial marker genes based on macroscopic inflammation classification. (E) Heatmap (left) and spline curve (right) illustrating dynamic expression of epithelial markers over time in DSS-treated mice (GSE22307). (F) Violin plots of selected gene markers in the DSS-treated mouse model from the same dataset. (G) Dot plot of epithelial marker expression in single-cell RNA-seq data from DSS-treated mice (GSE264408), categorized by normal, acute, and chronic colitis. (H) Representative immunofluorescence images of mouse small intestine sections from control and DSS-treated animals stained for OLFM4 (red) and DAPI (blue). Yellow arrows indicate crypts. Images represent three independent biological replicates. Scale bar = 100 µm.

To understand better the healthy SC/TA response to an initial signal of injury and compare it to IBD tissue response, we analysed the DSS-mouse model bulk-RNAseq^25^ (GSE22307), and the sc-RNAseq ^26^(GSE264408) data where EPCAM^+^ epithelial cells were extracted from the total cell population (Figure 1F). SC/TA and Wnt gene expression markers as *Lgr5*, *Sox9*, *Ephb2* and *Axin2* decreased over the course of inflammation, indicating a general decline within the intestinal tissue *in vivo* that are altered by an inflammatory stimulus. Lgr5^+^ stem cells tend to recover at day 4, reflecting an initial response to restore this pool in healthy epithelium. We identified an upregulation *of Clu* expression correlated with rising levels of S100 Calcium-Binding Protein A9 (*S100A9*), a biomarker commonly upregulated in inflammatory diseases^27^ (Figure 1E-G). Interestingly, Olfm4 increased at the protein level as inflammation intensified in the small intestine of DSS-treated mice (Figure 1H). These findings show how SC/TA pools balance regeneration and barrier integrity during inflammation in healthy and IBD tissue.

### Characterization of epithelial cell types in IBD vs healthy tissue using a 3D patient-derived organoid model

To investigate epithelial cell heterogeneity in IBD cells in absence of active inflammatory conditions, we derived 3D-intestinal epithelial organoids from IBD patient’s biopsies that retain the unique genetic and phenotypic characteristics of patient’s tissue that had been affected by chronic inflammation^28–30^ (Figure 1A). The expansion of these cells allowed us to create simplified *in vitro* IBD models that maintain active stem cell niches, and ongoing differentiation characteristic of non-inflamed (IBD normal) patients’ cells (Figure 2A). We performed bulk-RNAseq of healthy (hPDOs) derived from healthy individuals and IBD-PDOs derived from IBD patient’s tissue: IBD patient 89 (P89), P90 and P99. All the IBD-PDOs exhibited similar gene expression profiles, with P99 displaying a more pronounced inflammatory phenotype (Supplementary Figure 2A-C). Healthy and IBD-P99 organoids of early (passage 3) and late passages (passage 10) were sequenced to assess potential differences in molecular characteristics over time in culture. This approach allowed us to determine that passage number did not have a major influence on their gene expression profile (Figure 2B-I and S2A).

**Figure 2.**
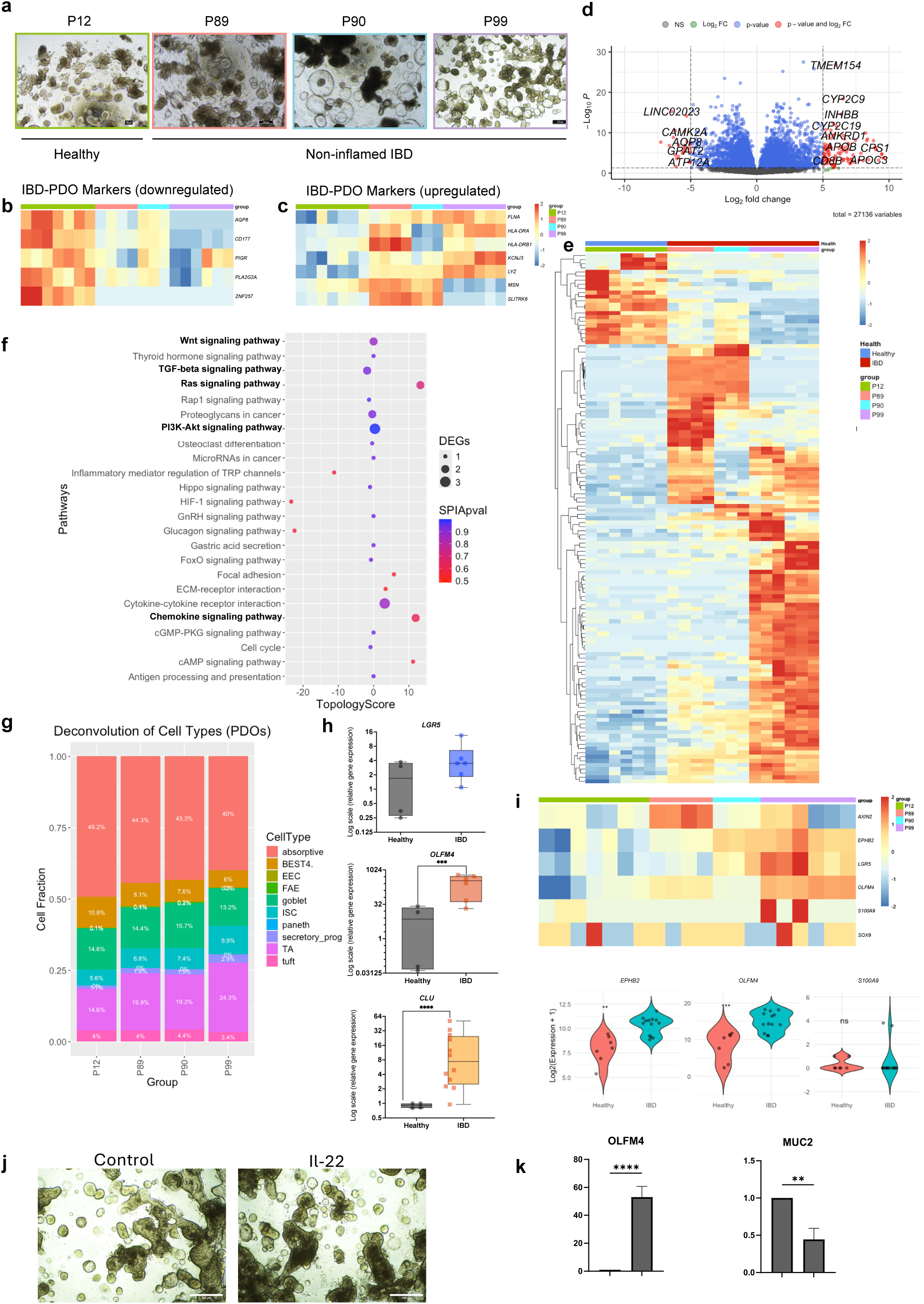
Characterization of epithelial cell alterations in IBD using a patient-derived organoid (PDO) model. (A) Representative brightfield microscopy images of healthy (hPDOs) and IBD-derived PDOs (IBD-PDOs). Scale bar = 100 µm. Bulk RNA-seq analysis of hPDOs (n = 7, results of 2 independent batches with early passage (n = 3, 3 independent wells) and late passage (n = 4, 4 independent wells) and IBD-PDOs (n = 13, results of 3 independent donors, P89 (n = 4, 4 independent wells), P90 (n = 3, 3 independent wells) and P99 (n = 6, results from 2 independent batches with early passage (n = 3, 3 independent wells) and late passage (n = 3, 3 independent wells). (B–C) Heatmaps showing expression of previously reported IBD-related markers (Dotti et al. 2017) in hPDOs and IBD-PDOs: (b) downregulated and (c) upregulated genes. (D) Volcano plot of differentially expressed genes (DEGs) between hPDOs and IBD-PDOs (adjusted *p* < 0.05, log₂ fold change > 5). Red dots represent significant DEGs; grey dots are non-significant. Total genes analysed: 26,771. (D) Heatmap of top DEGs distinguishing hPDOs from IBD-PDOs. (F) Bubble plot showing enriched pathways in IBD-PDOs vs hPDOs using Signaling Pathway Impact Analysis (SPIA). (G) Bar plot indicating estimated epithelial cell-type proportions in hPDOs and IBD-PDOs by computational deconvolution. (H) Bar plots showing mRNA expression levels of *LGR5*, *OLFM4*, and *CLU* in hPDOs (*n* = 4) and IBD-PDOs (*n* = 5). Results of 3 independent experiments. (I) Heatmap of selected epithelial gene markers in hPDOs and IBD-PDOs. (J) Brightfield images of hPDOs treated with IL-22 or vehicle control. Scale bar = 500 µm. Representative images from three independent biological replicates. (K) Bar graphs showing increased *OLFM4* and decreased *MUC2* mRNA expression in IL-22-treated hPDOs relative to control, as assessed by qPCR (*n* = 3) Results of 3 independent experiments.

The RNA-seq data was first validated by analysing previously reported IBD-PDOs biomarkers^31^: downregulated genes (*AQP8*, *CD177*, *PIGR*, *PLA2G2A*, and *ZNF257)* (Figure 2B) and upregulated genes (*FLNA*, *HLA-DRA*, *HLA-DRB1*, *KCNJ3*, *LYZ*, *MSN*, and *SLITRK6)* of IBD- vs hPDOs (Figure 2C). We identified 80 upregulated and 20 downregulated DEGs in IBD- vs hPDOs (fold change cutoff=5, p-value<0.05), where IBD-PDOs exhibited upregulation of genes involved in inflammatory pathways, such as cytokines and chemokines as IL-1β and IL-6 (Figure 2D, E). By Signalling Pathway Impact Analysis, we found other enriched pathways such as Wnt, PI3K-Akt and Ras, and downregulated pathways such as TGFβ signalling in IBD-PDOs vs hPDOs reflecting the impact of chronic inflammation on intestinal non-inflamed IBD epithelial tissue which included metabolic shifts, epithelial barrier function and, key signalling pathways involved in regeneration (Figure 2F and S2B).

To quantify and compare cell type content in both type of PDOs, we employed deconvolution analysis using CIBERSORTx with a signature matrix constructed using sc-RNAseq data from healthy human intestinal epithelial cells as reference^32^. We found that TA and Paneth cells were increased in IBD- compared to hPDOs while BEST4^+^ and enterocyte pools were decreased (Figure 2G). By qPCR we detected a significant increase of *OLFM4* in IBD-compared to hPDOs, however we found no significant change in *LGR5 and CLU* expression (Figure 2H and S2C). Moreover, the bulk-RNAseq gene expression analysis confirmed the significant upregulation of *OLFM4* and *EPHB2* (TA and Wnt target gene, respectively) in IBD-vs hPDOs (Figure 2I).

IL-22 is a cytokine that has a pivotal role in enhancing epithelial proliferation and stem cell-driven regeneration ^33,34^ which are essential processes disrupted during IBD. So, we analysed IL-22 pathway activation in human hPDOs to study OLFM4^+^ proliferative cells function in repair and regenerative intestinal mechanisms^35,36^. To that end we extracted EPCAM^+^ epithelial cells from the total population of sc-RNAseq data^35^ (GSE189423) and analysed the DEGs (Figure 2D-G). Interestingly, one of the top upregulated genes was *OLFM4* (Figure 2I) which we further validated by qPCR in our IL-22 treated vs control hPDOs. In parallel, IL-22 treatment reduced goblet cells detected by the decrease of *MUC2* expression marker (Figure 2J, K). Our findings identify OLFM4^+^ pool as key players of IBD epithelium and in epithelial repair mechanisms induced in healthy epithelium.

### Characterization of SC/TA and BEST4^+^ pools in healthy and IBD non-inflamed and inflamed tissue

We next analysed the heterogeneity of intestinal SC/TA and BEST4^+^ cell pools as these are key epithelial cells involved in regeneration by using publicly available sc-RNAseq data^15,37^. We extracted EPCAM^+^ epithelial cells from IBD non-inflamed (IBD normal) and inflamed tissues, and from healthy individual biopsies (UC=18, healthy=12 and, CD=46, healthy=25). Cell type annotations were standardized by using epithelial reference dataset (Figure 3A) (GSE185224) and labels of the cell types were transferred into the selected sc-RNAseq data using SingleR, ensuring unbiased labelling across datasets^38^ (Figure 3B and S3A). We then extracted SC/TA, and BEST4^+^ cell populations data, the latter being part of the enterocyte lineage directly derived from progenitors. We separated the data into healthy, non-inflamed (or normal), and inflamed groups, and observed a decrease in the SC/TA and BEST4⁺ pools in IBD inflamed tissue compared to healthy samples (Figure 3C and S3B).

**Figure 3.**
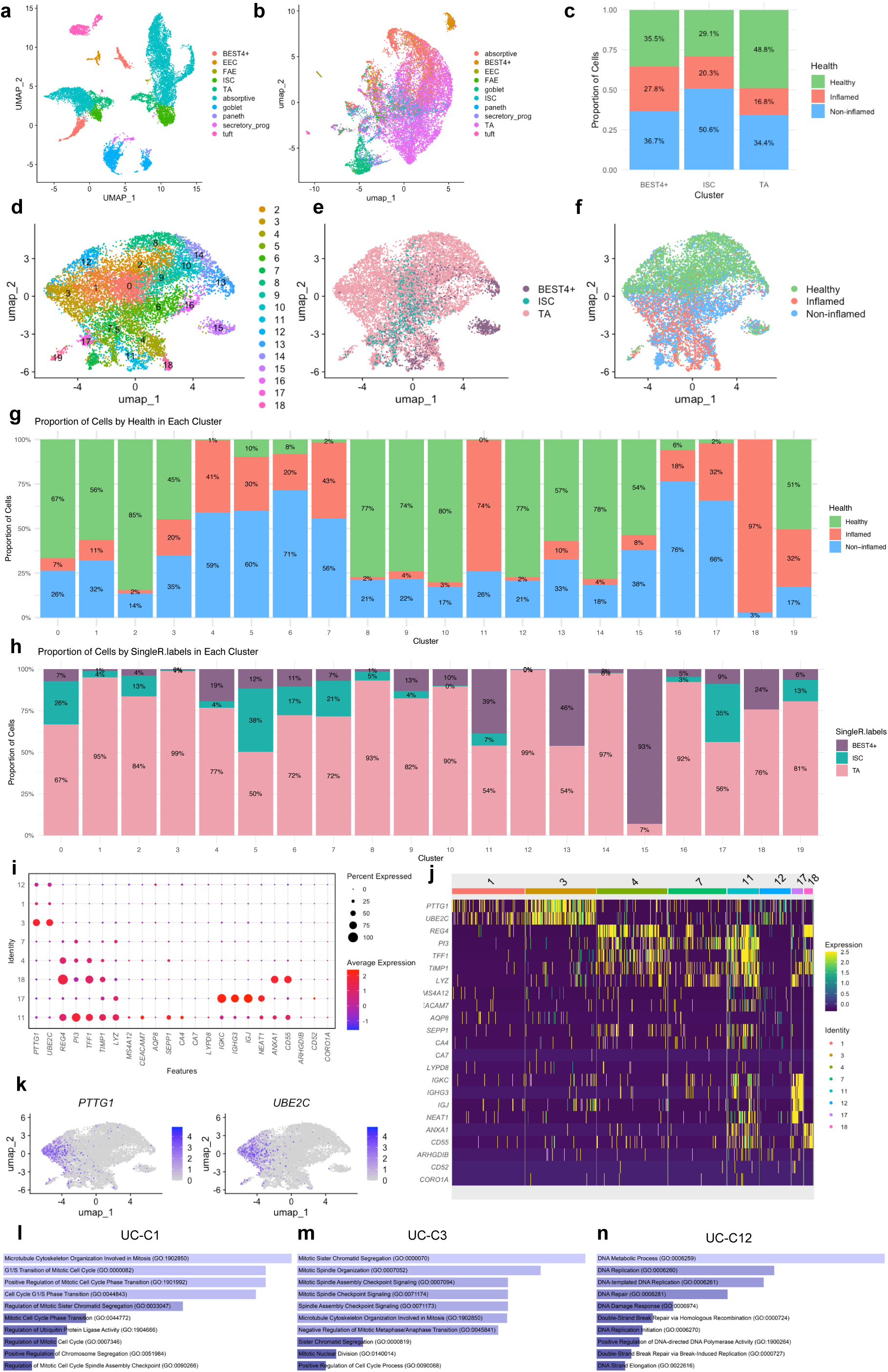
SC, TA and BEST4⁺ epithelial subpopulations in healthy, non-inflamed, and inflamed UC tissue. (A) UMAP embedding of epithelial cells from the reference dataset (Burclaff et al., 2022), annotated by cell type. (B) UMAP showing cell type predictions in the UC dataset (SCP259), highlighting SC/TA and BEST4⁺ epithelial subpopulations. (C) Bar plot quantifying SC/TA and BEST4⁺ cells across tissues with normal appearance, possible inflammation, and active inflammation. (D) UMAP displaying 19 transcriptionally distinct epithelial clusters (UC-C0 to UC-C18) identified in the UC dataset, indicating cellular heterogeneity within SC/TA and BEST4⁺ populations. (E–F) UMAPs annotated by (E) predicted cell type and (F) tissue health status. (G) Bar plot showing the distribution of UC clusters across epithelial cell types. (H) Bar plot showing the distribution of UC clusters by tissue health status. (I) Dot plot and (J) heatmap illustrating representative gene markers for biologically distinct epithelial clusters. (K) Feature plots showing expression of *PTTG1* and *UBE2C* in SC/TA and BEST4⁺ cells within the UC dataset (SCP259), demonstrating transcriptional diversity across disease states. (L) Bar plot showing enriched pathways from Gene Ontology (GO-Biological Processes 2023) for the top 50 genes in cluster UC-C1 (left), -C3 (middle) and –C12 (right).

Next, we performed principal component analysis on the variable features in a total of 11,268 cells for UC and 9,894 cells for CD datasets. We identified clusters using the Louvain algorithm with results visualised via Uniform Manifold Approximation and Projection (UMAP). This analysis resulted in 19 transcriptionally distinct UC clusters, and 13 CD clusters highlighting the functional heterogeneity and roles of SC/TA and BEST4^+^ cell types (Figure 3D and S3C). Clusters were further classified based on the cell type (BEST4^+^, SC and TA) (Figure 3E and S3D) and health status of the source tissue (healthy, non-inflamed and inflamed) (Figure 3F and S3E). The relative abundance of these clusters was evaluated based on cell types and health status (Figure 3G, H and S3F, G). We used the top 50 gene markers of each cluster to identify representative pathways, cell types and gene ontologies enrichment analyses (Table S1). Of all the computed clusters, UC-C1, −3, −4, −7, −11, −12, −17, −18 and CD-C1, −5, −8, −12, - 13, aligned clearly with known biological processes so we further refined the markers specific to these clusters (Figure 3I, J and S3H, I). Interestingly, we identified six gene clusters (UC-C1, −3, −12 and CD-C1, −5, −8) that predominantly consisted of TA cells (Figure 3G and S3F) highly enriched in MKI67^+^ proliferative states, showing high expression of proliferative markers like *PTTG1*, and *UBE2C* (Figure 3I-K and Table S1). The gene clusters UC-C1, −3 and CD-C1, −5 reflected enrichment of genes associated with cell division, M-phase of cell cycle, and microtubule cytoskeleton organization in mitosis. UC-C12 and CD-C8 were enriched in markers of S-phase, DNA synthesis and replication (Figure 3L and S3H).

Furthermore, we observed that IBD-PDOs vs hPDOs had higher expression levels of the top 50 proliferative markers, likely highlighting these cells’ role in adaptive proliferative mechanisms (Figure S3I). We further analysed the top 50 markers from these clusters in the DSS-mouse model (GSE22307) and observed that the MKI67^+^ cluster markers showed a biphasic proliferative response of healthy cells to cellular damage. This response was characterized by an initial decrease in proliferation, followed by a subsequent replenishment of proliferative cells, surpassing the original gene expression levels (Figure S3J, K). We further validated increased expression of *OLFM4* in IBD inflamed and non-inflamed vs healthy tissue in most of the clusters (Figure S3L). Together these findings highlight relevant and unique dynamics of these clusters in the SC/TA and BEST4^+^ pools during inflammation in healthy and IBD epithelium.

### An immune-enriched gene cluster present exclusively in SC/TA and BEST4^+^ pools of IBD inflamed tissues

We further investigated clusters UC-C18 and CD-C12 as these were predominantly composed of IBD cells and were absent in healthy tissue (Figure 3H and S3G). Enrichment analysis of the top 50 gene markers from each cluster showed their involvement in immune-related pathways such as IL-17, TNFα, NF-kB and Oncostatin M (Figure 4A and S4A). Notably, top marker genes such as *ANXA1*, *REG4*, *TFF1*, and *TIMP1,* previously linked to mucosal healing, epithelial restitution, and regeneration, were highly expressed within these clusters^39–42^. Markers for UC: *ANXA1*, *REG4*, *TFF1*, *CD55*, *TIMP1*, and *PI3; and* for CD: *PI3, IGGL5, REG4, DUOXA2, TIMP1, and CXCL1)* were chosen for further analysis as their expression was specific to UC-C18 and CD-C12, respectively (Figure 4B and S4B). These markers were present in SC, TA and BEST4^+^ cell types (Figure 4C, D and S4C, D). In SPARC IBD UC and CD data, the top genes of UC-C18 and CD-C12 were able to stratify patients according to their macroscopic appearance. Moreover, the average expression of these genes was significantly upregulated in inflammation and possible inflammation groups when compared to the normal group. Interestingly, this upregulation was also statistically significant in inflammation vs possible inflammation group (Figure 4E, F and S4E, F). We observed that the average gene expression of this signature increased with inflammation status from day 0 to 6 of DSS-treated mice (GSE22307) (Figure 4G and S4G). In IL-22 treated hPDOs sc-RNAseq data (GSE189423) we found that *ANXA1*, *CD55*, *PI3* and *TIMP1* were increased in IL-22 treated human organoids vs control and all of them were expressed in LGR5^+^ stem cells (Figure 4H, I). Hence, the UC-C18 and CD-C12 gene signature, represent inflammatory biomarkers of the epithelial compartment, and show promise in molecularly stratifying IBD patients based on their macroscopic appearance.

**Figure 4.**
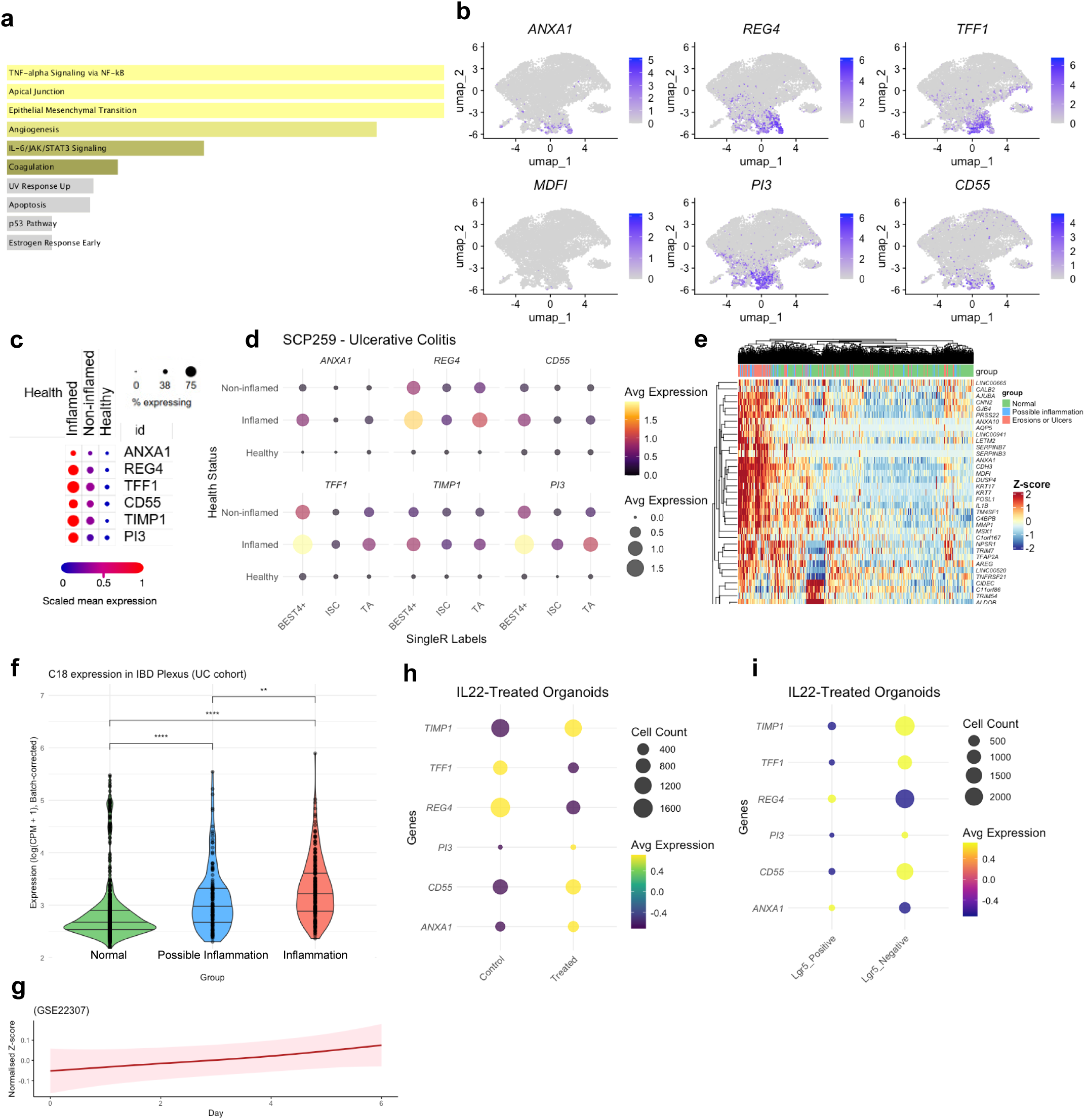
Immune-enriched gene clusters within SC, TA and BEST4⁺ epithelial populations in inflamed UC. (A) Bar plot showing enriched Hallmark gene sets from MSigDB for the top 50 genes in cluster UC-C18. (B) UMAP plots showing expression of selected UC-C18 marker genes (*ANXA1*, *REG4*, *TFF1*, *CD55*, *TIMP1*, and *PI3*) in SC/TA and BEST4⁺ cells from the UC dataset (SCP259). (C) Dot plot showing expression of the selected UC-C18 markers across epithelial cells stratified by tissue source: healthy, non-inflamed UC, and inflamed UC (SCP259). (D) Dot plot showing expression of the same UC-C18 markers across SC, TA, and BEST4⁺ epithelial subtypes, stratified by tissue condition. (E) Heatmap showing average expression of UC-C18 marker genes across SPARC IBD-UC epithelial samples, stratified by macroscopic appearance (normal, possible inflammation, and inflammation). (F) Violin plots showing UC-C18 gene expression patterns in SPARC IBD-UC samples grouped by macroscopic appearance. (G) Spline curve showing average expression dynamics of UC-C18 marker genes in DSS-treated mouse colitis dataset (GSE22307). (H) Dot plot showing UC-C18 marker expression in SC, TA, and BEST4⁺ populations from hPDOs treated with IL-22 or vehicle control (GSE189423). (I) Dot plot showing expression of UC-C18 markers stratified by LGR5 expression status in epithelial cells (GSE189423).

### Inflammation-Dependent Adaptive Dynamics of Epithelial Cells Categorized by a Novel Inflammation-Linked Gene Signature Associated with Disease Severity in IBD

We aimed to identify a reliable inflammation-associated gene signature to molecularly stratify patient samples according to the degree of inflammation and understand epithelial dependent inflammation dynamics. We curated an inflammation-related gene list (n=476) (Table S2) and compared their average gene expression in different clinical classifications of inflammation severity. We categorized the samples by clinical disease activity and found not significant differences between the groups defined by Mayo score for UC (mild, moderate, severe, and remission) and short Crohn’s Disease Activity Index (sCDAI) for CD (Figure 5A and S5A). However, when samples were categorized by macroscopic appearance recorded during endoscopy of IBD patients (normal or non-inflamed, possible inflammation, and inflammation) they formed three significantly distinct groups (Figure 5B and S5B). We then confirmed that *S100A9* gene expression was significantly upregulated in the inflamed samples when compared to possible inflammation or normal ones (Figure 5C and S5C). We further confirmed S100A9 protein upregulation in inflamed vs non-inflamed regions of IBD patient’s tissue samples (Figure 5D).

**Figure 5.**
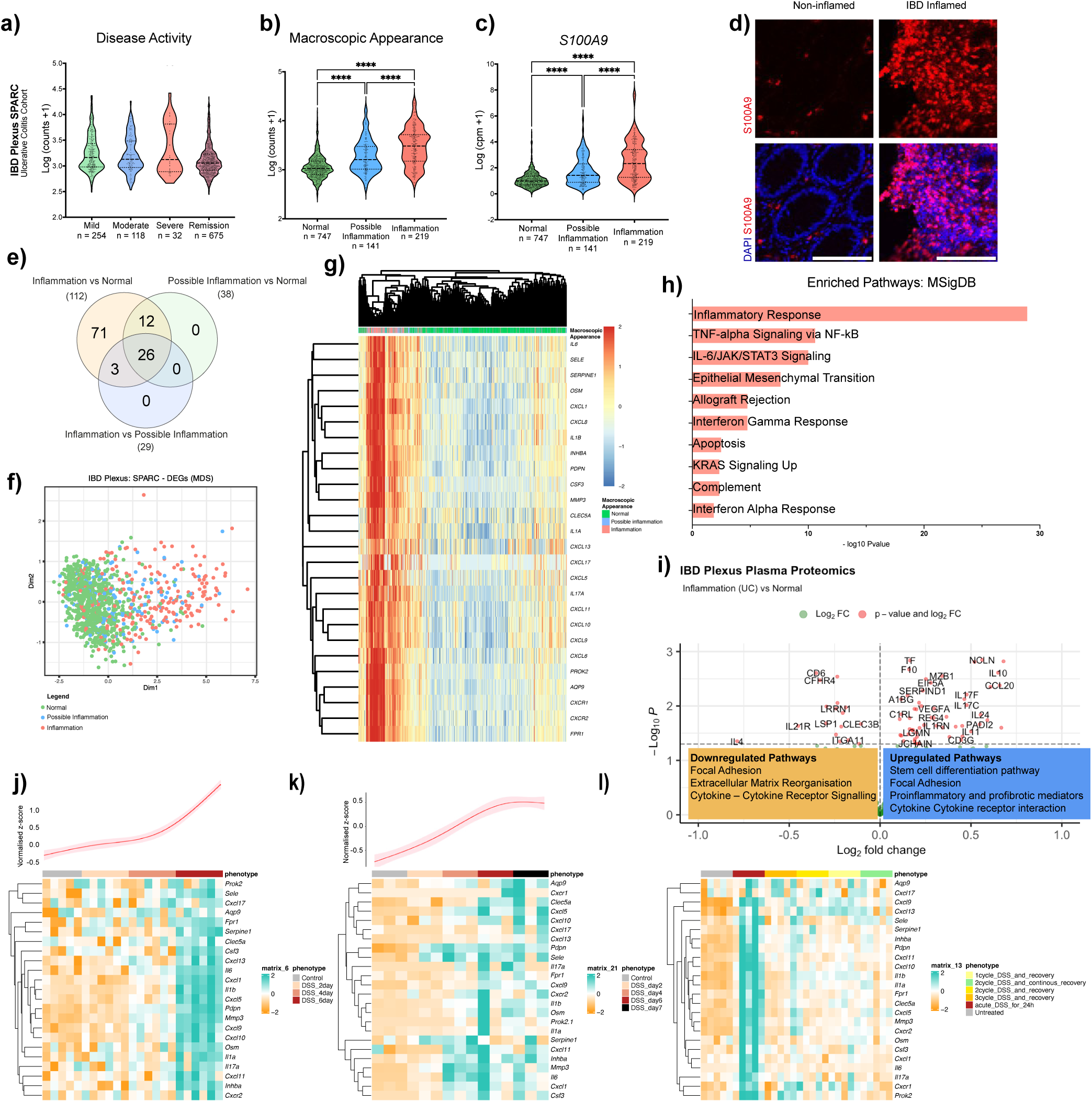
A novel inflammation-linked gene signature associated with inflammation severity in IBD. (A–C) Violin plots showing average expression of curated inflammatory genes (n = 476) in SPARC IBD-UC samples grouped by (A) disease activity, (B) macroscopic appearance, and (C) S100A9-based classification into normal (n = 747), possible inflammation (n = 141), and inflammation (n = 219) groups. (D) Representative confocal immunofluorescence images of S100A9 (red) and DAPI (blue) staining in inflamed and non-inflamed IBD tissues. Scale bar = 100 µm. (E) Venn diagram showing the intersection of significantly differentially expressed genes to define a 26-gene inflammation network in UC (26-GIN-UC). (F) Multidimensional scaling (MDS) plot showing sample distribution by gene expression profile across inflammation groups in the SPARC IBD-UC cohort. (G) Heatmap showing unsupervised clustering based on the 26-GIN-UC signature across the same cohort. (H) Significantly enriched pathways identified using Enrichr (MSigDB) from the 26-GIN-UC gene set. (I) Volcano plot showing differentially expressed proteins between inflamed and normal samples (red = significant proteins, adjusted *P* < 0.05; green = non-significant). Enriched upregulated (right) and downregulated (left) pathways are annotated from STRING analysis. (J–L) Heatmaps and spline curves showing dynamic expression of the 26-GIN-UC signature in DSS-treated mice from datasets: (J) GSE22307, (K) GSE214600, (L) GSE42768. P < 0.01 (**), P < 0.0001 (****).

We validated the categorization of these groups by using an inflammation-related gene signature (n=42) from a prior UC study^15^ (Figure S5D, E). In addition to the expected upregulation of inflammatory genes in the inflamed vs normal group, we found that possible inflammation category also showed elevated expression of inflammatory markers compared to the normal group suggesting an underlying genetic predisposition to inflammation that is not clear in tissue macroscopic assessments. Overall, we found that the macroscopic appearance is a reliable classifier of inflammation severity, hence, it was maintained for further analyses.

We then performed differential gene expression analysis to compare the inflammatory profile across these groups using the curated inflammation-related gene signature (n=478). In UC, we found 112 differentially expressed genes (DEGs) between inflammation vs normal samples, 38 DEGs between possible inflammation vs normal samples, and 29 DEGs between inflammation vs possible inflammation samples (Table S3). A Venn diagram illustrated that 26 genes (26-GIN-UC) were consistently differentially expressed across all three comparisons (Figure 5E). In addition, multiple dimensionality (MDS) analysis showed that this gene signature was effective in clustering the samples according to the clinical classifier of their macroscopic appearance highlighting their potential in distinguishing inflammation (Figure 5F). In CD, we found 380 DEGs between inflammation vs normal samples, 304 DEGs between possible inflammation vs normal samples and 35 DEGs between inflammation vs possible inflammation samples A Venn diagram illustrated those 30 genes (30-GIN-CD) shared among the three comparisons and MDS plots demonstrated effective sample stratification (Table S3 and Figure S5F, G). To further evaluate both gene signatures (26-GIN-UC and 30-GIN-CD), we performed unsupervised hierarchical clustering analysis on the SPARC IBD cohort which aligned with macroscopic appearance effectively clustering the samples according to the inflammation category. Enrichment analysis of these two gene signatures showed strong representation of inflammatory response pathways as TNFα and IL-6/JAK/STAT (Figure 5G, H and S5H, I). Comparison of inflamed vs normal samples from the proteomics SPARC IBD dataset (UC and CD) confirmed that members of inflammatory pathways such as interleukins IL-10, −24, −17 were amongst the significantly differentially expressed proteins (*p*-value < 0.05) (Figure 5I and S5J). Protein-protein interaction analysis showed that proteins related to focal adhesion, extracellular matrix reorganisation were downregulated, whereas proteins related to pro-inflammatory and profibrotic mediators were upregulated (data not shown).

We further validated these gene signatures in three different transcriptomic datasets of dextran sulphate sodium (DSS)-treated UC mouse model that tracks inflammatory responses and disease progression over time^43^. We identified the upregulation of the 26-GIN-UC and 30-GIN-CD signatures with increasing inflammation, observing the onset, peak, and resolution of inflammation in a time-dependent manner (Figure 5J-L, and S5K).

We then evaluated the effectiveness of the new identified gene signatures in stratifying IBD samples according to inflammation degree. First, we performed unsupervised hierarchical clustering on SPARC IBD UC samples, identifying five distinct groups based on their UC-C18 gene expression profiles. This clustering reflected the underlying biological patterns (endoscopic assessment/macroscopic appearance) thereby providing a novel unbiased approach for IBD samples stratification. We found that these groups corresponded to varying inflammation potential grades, ranging from 1 to 5, with 1 representing low inflammation and 5 representing severe inflammation. We validated this classification with the average expression of the 26-GIN-UC (Figure 5) and UC-C18 gene signatures, which showed a progressive increase with the degree of inflammation, confirming the appropriate stratification of the samples (Figure 6A-C). Notably, *S100A9*, showed a significant upregulation in the most inflamed samples, reinforcing the robust and unbiased framework for classifying IBD samples based on inflammation degree (Figure 6D).

**Figure 6.**
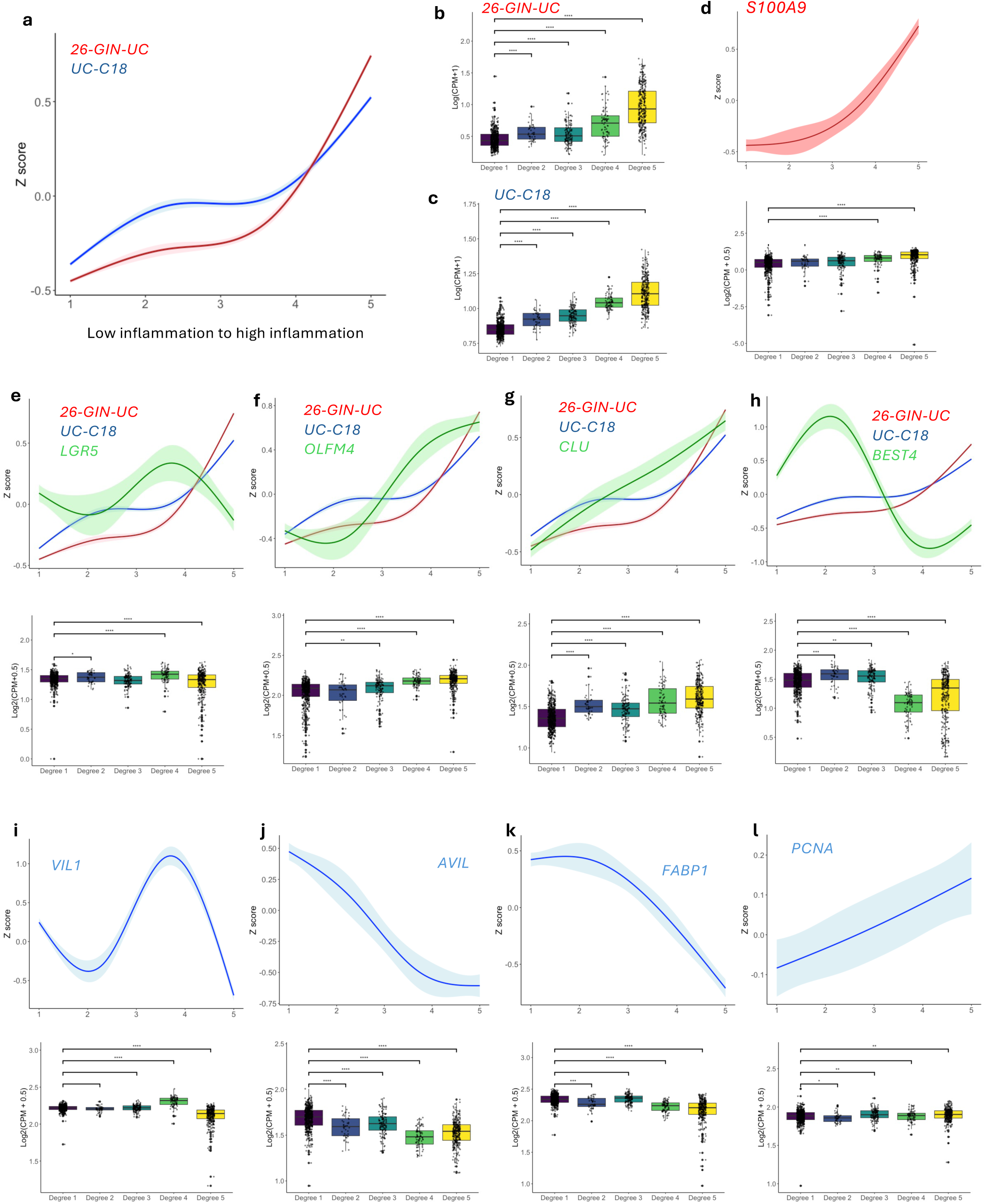
Adaptive epithelial responses in UC stratified by degree of inflammation. (A) Spline curve showing the average expression of the 26-GIN-UC and UC-C18 gene signatures across increasing inflammation scores. (B–C) Box plots showing expression levels of (B) 26-GIN-UC and (C) UC-C18 gene signatures across low to high inflammation categories. (D) Spline curve (top) and box plot (bottom) of *S100A9* expression stratified by inflammation severity. (E–H) Spline curves (top) and box plots (bottom) showing gene expression across inflammation severity for (E) *LGR5*, (F) *OLFM4*, (G) *CLU*, and (H) *BEST4*. (I–L) Spline curves (top) and box plots (bottom) for epithelial differentiation and proliferation markers (I) *VIL1*, (J) *AVIL*, (K) *FABP1*, and (L) *PCNA* across inflammation categories.

We overlaid the average expression of specific SC/TA and differentiated cell markers with both inflammatory signatures. *LGR5* expression, showed varying levels in different inflammation degrees, with an overall decrease in the highest inflammation degree 5 and weak negative correlation with (*r =* −0.085) with UC-C18 signature. The spline curve model shows the dynamic shift in *LGR5* expression in the different inflammation degrees (Figure 6E and S6). *OLFM4* expression exhibited a subsequent upregulation in degrees 3 to 5 with an overall positive correlation (*r =* 0.579) with the UC-18 signature (Figure 6F and S6). Similarly, the revival stem cell marker *CLU* increased with severity of inflammation with positive correlation (*r* = 0.542) (Figure 6G and S6), while *BEST4* was upregulated in degree 2 but its expression gradually decreased in degree 3 and 4 (Figure 6H). *VIL1*, gene expressed in enterocytes lining the intestinal villi and crypts, showed a dynamic expression from degree 2 to degree 4 (Figure 6I). *AVIL*, gene marker for brush border integrity and actin dynamics and *FABP1* enterocyte marker were both decreased upon severity of inflammation (Figure 6J, K). We observed that proliferation markers such as *PCNA* exhibited a general increase when compared degree 5 vs degree 1 (Figure 6H) that reflects the balance between inflammation-induced epithelial injury and the subsequent regenerative response. Moreover, we observed a similar trend of negative correlation of *BEST4*, *VIL1*, *AVIL* and *FABP1* with UC-18 expression (*r* = −0.3, −0.35, −0.536, - 0.496, respectively). (Figure S6). We obtained very similar results for CD dynamic shifts (CD-C12, and 30-GIN-CD) (Figure S7). We evaluated the robustness of these signature using patient level hold out test our models performed very well for “Normal” samples, while classification of inflamed states was more modest and the intermediate “Possible inflammation” cases were not well predicted as it likely represents a heterogeneous mix of early, partial, or ambiguous lesions, and may also capture endoscopic uncertainty (Figure S7p,q). To address this, we also applied unsupervised clustering of the gene signature followed by dendrogram-based cutting to derive an alternative, data-driven ordering (Figure 6). This approach produced a molecularly defined progression of five groups that may better reflect underlying biology. This complementary analysis highlights that transcriptomic ordering can resolve heterogeneity within inflamed states. In addition, we also evaluated whether clinical parameters had an effect on the ability of this gene signature to stratify inflammation. We found that the gene signature score remained strongly associated with macroscopic appearance after adjusting for sex, race, and smoking status, with direction and magnitude consistent across models for both UC and CD. None of the covariates materially altered the effect of the signature, indicating that the association is independent of demographic or smoking-related factors (Supplementary Table S5, S6).

The combination of scatter plots and spline curve model provided a comprehensive approach in understanding how the average expression of UC-C18 and CD-C12 gene signature reflects dynamic changes in different cell type markers. While scatter plots offer a more rigid, point-by-point correlation, spline curves capture smoother, more continuous transitions in expression patterns within our patient stratifications. Together, these methods show inflammation-dependent SC/TA and BEST4^+^ cells dynamics and demonstrate that these pools derived UC-C18 and CD-C12 gene signatures reflect both broad and subtle dynamic shifts across different inflammation degrees.

### Exploring molecular mechanisms in IBD by data integration of UC and CD sc-RNAseq datasets

We then performed integration of the epithelial cell compartments of SCP259 (UC) and SC1884 (CD) to ensure comparability across both IBD subtypes, UC and CD. This revealed overlapping epithelial cell populations (Figure S8A-D). We applied standardised automated cell annotations with CellTypist, referencing the pan-gastrointestinal (pan-GI) atlas, with minor manual adjustments (Figure S8E). This analysis revealed a marked depletion of colonocytes in inflamed samples, accompanied by a notable expansion of deep crypt secretory cells (DCS), which are thought to represent the colonic equivalent of small intestinal Paneth cells (Figure S8F).

To confirm the combined changes associated with inflammation in both IBD subtypes, we performed unsupervised clustering of gene expression profiles based on disease status, grouping inflamed UC and CD samples into “inflamed”, “non-inflamed” category and “healthy” samples (Figure S8G). This analysis identified a core set of 20 top differentially expressed genes that were consistently upregulated in inflamed UC/CD but exhibited minimal expression in non-inflamed and healthy samples (Figure S8H). We used these genes to create an inflammatory gene score to assess cell-type-specific expression patterns, which revealed that they are predominantly expressed in the DCS population, with limited expression across other cell types (Figure S8I). Transcription factor activity mapping across various cell populations further identified NFκB activation in DCS as a key feature of inflamed state, consistent with findings from the differential expression analyses comparing IBD inflamed vs non-inflamed cells in DCS and stem cells (Figure 7A-D). Trajectory analysis indicated that the differentiation trajectory of intestinal SCs is disrupted during inflammation, with a skewing toward DCS lineages (Figure 7E). Genes related to DCS trajectory were expressed higher in inflamed tissue relative to the healthy samples (Figure 7F).

**Figure 7.**
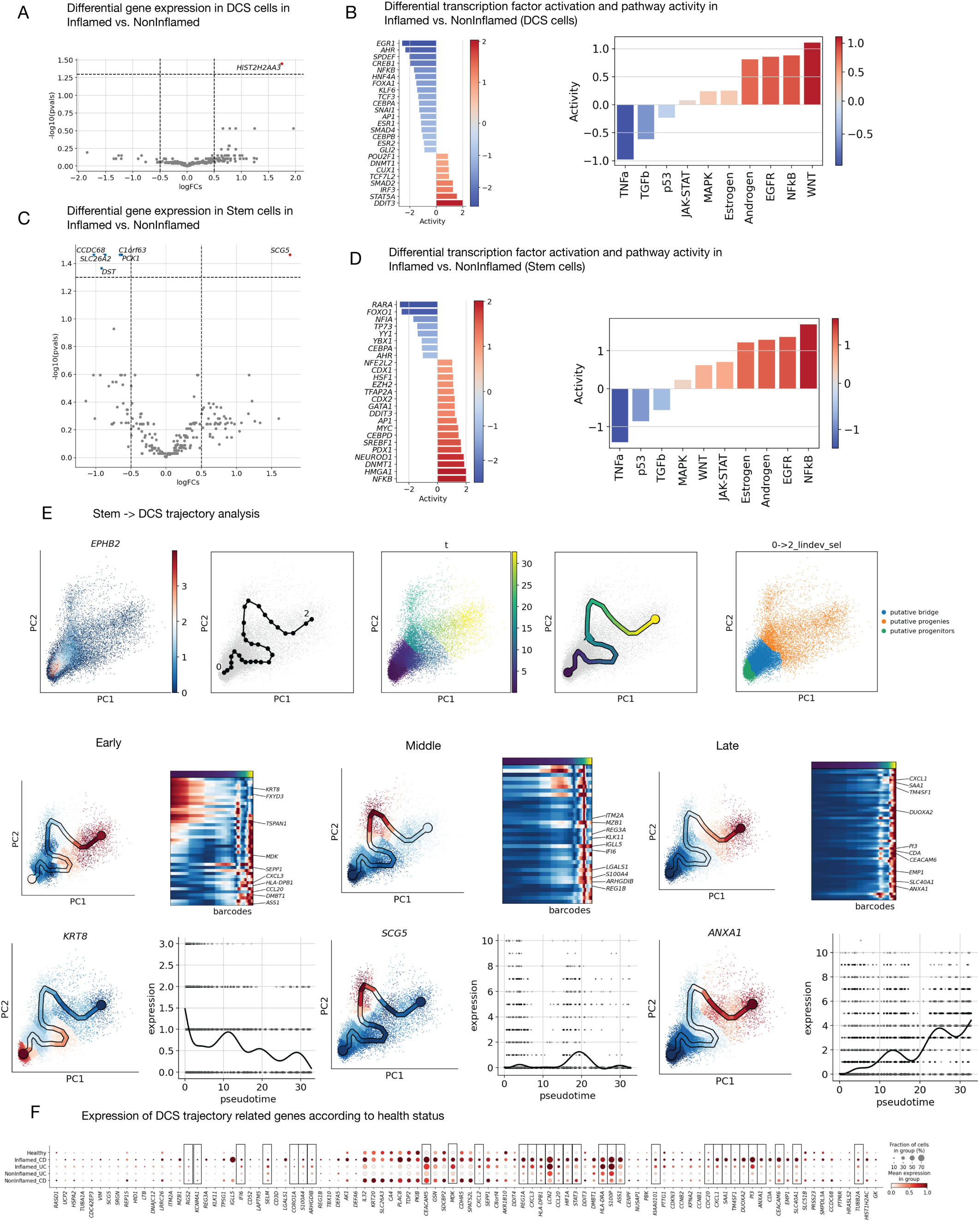
Integration of UC and CD sc-RNAseq data reveals distinct epithelial mechanisms of inflammation. Differential gene expression and transcription factor pathway activation in DCS (A, B) and SCs (C, D) in Inflamed vs non-Inflamed groups. (E) Basic trajectory pseudotime analysis of the route from SCs to DCS. Taking the middle and late expressed genes in this trajectory (*KRT8*-early, *SCG5*-middle and *ANXA1*-late). (F) Bubble plot showing the expression of DCS trajectory related genes according to health status.

Together, these findings suggest that inflammation disrupts normal epithelial differentiation, driving SCs toward a DCS-like fate characterized by expression of pro-inflammatory genes. Notably, the data identifies biomarkers of inflammation-associated stem cell reprogramming in the colonic epithelium.

## Discussion

Accurately evaluating inflammation severity in IBD remains a clinical challenge. Existing approaches such as symptom-based scores and biomarker assessments, frequently lack the sensitivity and specificity needed to reliably reflect the extent of inflammation. Endoscopy, often followed by histological examination, remains the gold standard but is invasive, costly, and not readily accessible in all settings. This underscores the urgent need for robust, non-invasive, and reproducible molecular methods to assess inflammation in IBD. Such tools are crucial for improving patient stratification, guiding treatment decisions, and deepening our understanding of epithelial and immune-driven tissue pathology^44–46^. Traditional clinical indices obtained by endoscopies, such as the sCDAI for CD and the Mayo score for UC, are widely used but often fail to reflect the molecular and cellular heterogeneity of inflammation observed at the tissue level^47,48^. In this study, we found that macroscopic appearance assessed during endoscopy closely aligned with inflammation-associated gene expression, suggesting its possible future potential to serve as a biologically meaningful classifier of inflammation severity rather than overall disease activity. Through differential gene expression analysis based on macroscopic appearance, we identified distinct inflammation-associated gene signatures for UC and CD. These signatures offer an unbiased approach for stratifying patient samples and provide valuable insights into epithelial cell diversity, regenerative responses, and barrier function across varying degrees of inflammation in IBD disease subtypes. These findings are preliminary and require prospective validation before they can be proposed as clinical tools.

Understanding how different epithelial cell populations, including stem and progenitor cells, react to varying levels of inflammation is essential for unravelling the processes of mucosal injury and healing. Our comprehensive profiling of epithelial cell heterogeneity across different inflammation stages reveals significant and coordinated shifts in the different epithelial cell pools. Through a multimodal analytical approach, we observed a reduction in TA cells as inflammation severity increased, a trend previously associated with impaired proliferative capacity during chronic inflammation or skewed differentiation dynamics^16,49^. In contrast, we observed that LGR5^+^ stem cells varied across inflammation duration and intensity, which possibly suggests a constant compensatory mechanism aimed at restoring epithelial barrier integrity. Interestingly, OLFM4⁺ proliferative cells exhibited sustained upregulation during inflammation, reflecting a regenerative response to counterbalance epithelial damage, and this expression remained elevated even in non-inflamed IBD tissue further highlighting the expansion of proliferative stem cells as a hallmark of IBD. Our findings revealed that these dynamics were substantiated in mouse healthy tissue^50^, where inflammation led to the decrease of TA pool along with an increase in Clu⁺ and Olfm4⁺ stem-like populations. This new identified epithelial functional plasticity driven by inflammation has potential implications for barrier dysfunction and regeneration.

Under inflammatory conditions, the proliferation of SC/TA cells is predominantly regulated by pathways such as Wnt/β-catenin and Ras signalling. These pathways are essential for sustaining the equilibrium between self-renewal and differentiation within the stem cell niche^6,10,51,52^. Inflammatory cytokines such as TNFα and IL-6 are known to enhance the proliferation of intestinal stem cells by activating signalling pathways like STAT3. This activation can amplify the inflammatory milieu and contribute to an excessive proliferative response within the tissue^53–56^. Our mechanistic insights and SC/TA dynamics data indicate that while enhanced proliferation may initially facilitate tissue repair, it can also lead to dysregulation in chronic inflammatory settings. This may result in depletion of the stem cell reservoir or inappropriate differentiation, ultimately weakening the intestinal barrier and contributing to the sustained epithelial injury seen in IBD^36^. Targeting specific proliferation-related pathways within this epithelial clusters may offer therapeutic potential by fine-tuning regenerative responses to improve barrier integrity while avoiding the overactivation that contributes to disease pathology.

We found that IBD tissue classified as non-inflamed still exhibits subtle transcriptional and cellular alterations compared to healthy control, suggesting a baseline state of immune activation and epithelial stress even in macroscopically normal regions^16,57^. To explore epithelial cell-type alterations between IBD and healthy tissue in the absence of overt inflammation, we leveraged 3D PDO models, which we confirmed that it recapitulates the cellular composition and regenerative dynamics of the intestinal epithelium^29–31^. IBD-PDOs displayed a distinct epithelial profile compared to hPDOs, including upregulation of inflammatory mediators and SC/TA proliferative markers such as OLFM4. These changes were accompanied by a reduction in differentiated epithelial cell types, such as enterocytes and BEST4⁺ cells, and an increase in Paneth cell populations, suggesting a shift toward a regenerative or stress-associated state. This reinforces the concept that epithelial cells in IBD retain a molecular imprint of inflammation, even in histologically normal areas^16,58,59^. Additionally, IL-22 pathway activation in hPDOs showed that OLFM4⁺ cells are activated during epithelial repair, supporting their role as mediators of regeneration in both homeostatic and disease settings. We highlight the utility of PDO models to dissect functionally epithelial alterations in IBD offering insights into therapeutic response.

We further characterized SC/TA and BEST4^+^ epithelial clusters uniquely enriched in inflamed IBD tissues but absent in healthy samples. These clusters were significantly enriched for immune-related pathways that are well-established drivers of chronic intestinal inflammation and epithelial remodelling in IBD^60,61^. This enrichment suggests the emergence of inflammation-specific epithelial transcriptional states during active disease. We report here, for the first time, an inflammation-associated epithelial gene signature specific to stem and progenitor cells. This signature successfully stratified patients based on their macroscopic inflammation degree in both UC and CD. The observed enrichment of SC/TA signatures in inflamed tissue aligns with earlier reports of crypt hyperplasia and stem cell activation as hallmark features of IBD pathology^56,62^. Thus, these transcriptional states likely represent inflammation-induced epithelial remodelling programs, offering a refined molecular framework for patient stratification and potential therapeutic targeting of regenerative responses in IBD.

Our data suggest that inflammation alters the differentiation trajectory of intestinal stem cells, skewing them toward DCS cell lineages, which are implicated in epithelial barrier maintenance and immune regulation ^16^. This shift occurred alongside a relative depletion of colonocyte differentiation and was also evident in non-inflamed IBD-PDOs when compared to healthy controls. This adaptability can be crucial in chronic inflammatory conditions as stem cells must maintain tissue integrity despite continuous stress. However, it also raises questions about long-term consequences, as chronic inflammation can lead to exhaustion or altered differentiation patterns in stem cells, potentially impacting their regenerative capabilities over time^63,64^. These findings reflect a broader epithelial remodelling driven by disrupted lineage commitment. Notably, we identified a core set of inflammation-associated genes highly expressed in DCS, with NF-κB signalling emerging as a prominent regulator that requires direct experimental follow-up for functional validation. This aligns with previous studies highlighting the central role of the NF-κB pathway in mediating epithelial responses after injury and sustaining chronic inflammation^65,66^. Furthermore, NF-κB activation has been shown to modulate Wnt pathway in colonic mouse tissue and specific ablation of RelA/p65 retards crypt stem cell and progenitors expansion^67,68^. Future investigations may help to determine whether modulating this pathway in DCS could lead to new therapeutic opportunities for IBD. However, selectively targeting NF-κB signalling in DCS is technically challenging due to the complex cellular composition of the intestinal mucosa, intestinal region-specific identity, and the broad physiological roles of NF-κB in maintaining epithelial integrity and immune balance^65,66,69^. Furthermore, several studies have demonstrated the complex interactions between individual NF-κB subunits in regulating colonic susceptibility to inflammation^69–71^. In addition, off-target modulation could disrupt homeostatic mechanisms, potentially leading to adverse consequences such as impaired wound healing, or dysregulated inflammatory responses. Overcoming these barriers will require advances in targeted delivery systems, and rigorous preclinical validation to ensure safety and specificity for future therapeutic applications.

In summary, our findings highlight how inflammation-induced alterations in epithelial stemness, and differentiation contribute to the dysfunction of the intestinal barrier in IBD. The capacity of stem and progenitor cells to adapt under inflammatory stress offers valuable insights for enhancing tissue repair strategies. Targeting these adaptive mechanisms may support the development of novel therapies that sustain epithelial regeneration, minimize disease flares, and foster long-term remission. Understanding the link between epithelial plasticity and specific inflammatory cues may ultimately guide precision treatments aimed at improving clinical outcomes and patient well-being.

## Methods

### Ethical Review

The research complies with all relevant ethical regulations and guidelines. Intestinal biopsy samples were obtained from the Queen’s Medical Centre of Nottingham (UK) with patient consent from IBD patients and histologically healthy individuals in accordance with ethical guidelines (17/EM/0126).

Animal DSS-experiments were performed in accordance with protocols approved by the Service de la Consommation et des Affaires Vétérinaires of Canton Vaud, Switzerland (VD1052/VD1053). The work has been reported in line with the ARRIVE guidelines 2.0. Section expanded in the “Declarations” section.

### IBD Plexus Cohort

The patient cohort included in this study was derived from the Study of a Prospective Adult Research Cohort with IBD (IBD Plexus Study of a Prospective Adult Research Cohort, SPARC IBD) (UC=1,097, CD=2,138)), a part of the Crohn’s & Colitis Foundation IBD Plexus program^21^. This includes a geographically diverse research cohort of patients with IBD using standardized data and biosample collection methods and processing techniques. We included patients from the SPARC IBD cohort in our analysis, who had a history of UC and CD and had available bulk-RNAseq of biopsies. We also included plasma proteomics, where applicable, for patients within the cohort. SCDAI for CD and 6-point MAYO for UC patients scores were calculated for patients with sufficient data at enrolment and assigned a disease severity classifier:

**Table 1.** SCDAI for CD and 6-point MAYO for UC patients scores for disease activity classification.

| Disease Activity | sCDAI | 6-point MAYO |
| --- | --- | --- |
| Remission | <150 | 0-1 |
| Mild | 150 – 219 | 2-3 |
| Moderate | 220-450 | 4-5 |
| Severe | >450 | 6 |

### Data acquisition for bioinformatic analysis

All the data processing and analysis was done in RStudio (Version 4.2.0 and 4.3.0) or Python.

#### IBD Plexus data

The raw counts of bulk-RNAseq of UC and CD patients derived from biopsy samples within the SPARC IBD cohort were batch corrected using *Combat_seq*, followed by normalisation and log transformation. Multi-dimensionality Scaling (MDS) analysis was performed to check the sample distribution. SPARC IBD samples were compared against different levels of histological inflammation: erosions or ulcers (or inflammation) vs normal, erosions or ulcers vs possible inflammation (or non-inflamed) and erosions or ulcers vs possible inflammation, for a curated inflammatory gene list (n=476) (Supplementary Table 2). Differential gene expression analysis for the different groups in IBD Plexus Cohort was conducted using *edgeR*^72^ and *limma*^73^. P-values threshold was set at 0.05 and adjusted using the Benjamini-Hochberg method.

#### Human sc-RNAseq data

Pre-processed and annotated sc-RNAseq data containing EPCAM^+^ epithelial cells from human tissue was obtained from Single Cell Portal of the accession numbers SCP259, SCP1884 and the reference dataset from Gene Expression Omnibus (GEO) under the accession number GSE185224. SCP259 contained cells from both non-inflamed and inflamed tissues data from UC, and healthy individuals’ biopsies (UC=18, healthy=12). SCP1884 was a comparable dataset with CD patients and normal individuals’ biopsies (CD=46, healthy=25). Cells with mitochondrial gene content exceeding 5%, features fewer than 200 and more than 2,500 were excluded to retain the highest quality cells. Each dataset was processed according to standard workflow using *Seurat*^74^ (version 5, package in R. Cell annotation was uniformly performed using SingleR to predict labels using the reference dataset. The counts matrix from both the datasets was used as the query, while the GSE185224 reference counts were used for the reference labels. Cell type labels were predicted using the Wilcoxon differential expression method. The predicted labels were added to the Seurat object and was subset to include only cells with “TA”, “BEST4^+^” and “SC” labels and this was used for further analysis.

For single-cell dataset integration studies, SCP259 and SCP1884 raw matrices were imported using scanpy (v1.10.4) and cells were filtered to exclude cells with greater than 20% mitochondrial counts and greater than 6000 features. Next, cells were processed through scrublet and predicted doublets excluded. Counts per cell were normalised to a target of 1×10^4^ and then log-transformed. Highly variable genes were selected followed by principle-component analysis (PCA), nearest-neighbour estimation (n_neighbours = 5, n_pcs = 17, knn=True) and UMAP projection followed by leiden clustering (resolution, 0.3). To correct for batch-effects across the two datasets, batch-balanced k-nearest neighbours (bbknn) correction was performed using the ‘Subject’ as batch key and leiden cluster as confounder. PCA, nearest neighbour, UMAP projection and leiden clustering were repeated using the same settings. Automated cell-type annotation was performed using CellTypist majority voting using the Pan-GI cell atlas model (:https://cellgeni.cog.sanger.ac.uk/gutcellatlas/celltypist_models/2_full_healthy_reference_AP_all_organs_finalmodel.pkl). Each leiden cluster was annotated with the predicted cell label. To check the validity of annotation, cluster specific gene expression was ranked, using the Wilcoxon method. In most cases, marker genes accorded with known markers of each subset, but in a few cases (e.g. Stem_2), gene expression suggested misclassification (e.g. Paneth cell) and therefore these labels were manually corrected. In addition, some clusters were merged (e.g. Goblet_1, Goblet_2) where leiden clustering had separated cells of the same type. Trajectory analysis of specific subsets was performed using scfates and decoupler was used for pseudobulk analysis of cell populations across health states to determine differential gene expression and transcription factor activity. Finally, genemania (genemania.org) was used for network and gene-set functional analysis of differentially expressed genes.

#### Cluster UC-C18 and CD-C12

The subset data with progenitors’ cells for each dataset were handled individually and processed according to standard workflow from the Seurat package in R. Dimensionality reduction and Louvain Leiden clustering (resolution 0.1 −1.5) were carried out and cell lineages were annotated based on enrichment analysis and defined marker gene expression of each cluster.

#### DSS-treated mouse models sc-RNAseq data

Single-cell RNAseq data was obtained from GSE264408 was utilised to study the cell type changes in normal, acute and chronic mouse models. Raw matrices were loaded for each sample using the Seurat package. Each sample was processed into a Seurat object, applying filters for quality control: cells with 200–4,000 detected genes and mitochondrial content <10% were retained. Metadata including sample identifiers and condition types were added to Seurat objects. Seurat objects for all samples were merged into a single dataset and standard workflow was applied. Data were further harmonized using the Harmony algorithm to correct batch effects, and integrated clusters were identified using shared nearest neighbour graphs and clustering algorithms. To focus on epithelial S/P cells, Epcam^+^ cells were subset (Epcam > 0.5). Dimensionality reduction (UMAP) and clustering were repeated. Epcam^+^ cells were categorized based on “Lgr5” expression into “Lgr5_Positive” and “Lgr5_Negative” groups. Wilcoxon tests assessed differential expression significance across conditions. Visualizations included violin plots and feature plots for key markers.

#### DSS--treated mouse models microarray transcriptomic data

Raw expression data of GSE22307 and GSE42768 was downloaded from NCBI-GEO platform and log2 normalised using limma package. Similarly, GSE131032 and GSE214600 raw counts data downloaded from same platform. To process raw counts, we used deseq function of Deseq2 package. Z-score was calculated from normalised data to visualize gene expression in either heatmap or spline curve using pheatmap and ggplot2.

#### PDOs sc-RNAseq data

Raw sc-RNAseq of IL-22 treated and untreated intestinal human organoids were retrieved GSE189423, merged, and batch effects were corrected using Harmony integration. EPCAM^+^ cells were identified and further analyzed through dimensionality reduction (UMAP), clustering, and marker gene expression profiling. To focus on epithelial stem/progenitor cells, Epcam^+^ cells were subset (Epcam > 0.5). UMAP and clustering were repeated. Epcam^+^ cells were categorized based on “Lgr5” expression into “Lgr5_Positive” and “Lgr5_Negative” groups. Wilcoxon tests assessed differential expression significance across conditions. Visualizations included violin plots and feature plots for *ANXA1* and *REG4*.

#### PDOs bulk-RNAseq data

RNA collected from healthy (P12) and IBD organoids (P89, P90, P99) and their mRNA was sequenced. A custom pipeline with standard bulk-RNAseq workflow was implemented for quality control, trimming and alignment. Aligned reads were quantified using FeatureCounts.

### Pathway Enrichment Analysis

Key pathways of the DEGs were performed in MSigDB database using the web tool, Enrichr. Pathways enriched for the differentially expressed proteins in plasma proteomics of SPARC IBD cohort was identified using KEGG database using protein-protein-interaction networking using the web tool, StringDB. Gene Ontology analysis for the cluster makers was performed using web tool, Enrichr. DEGs with an unadjusted p-value < 0.05 were considered significant. For downstream pathway analysis, we further filtered for DEGs with an adjusted p-value (padj) < 0.05 and an absolute log2 fold change > 5. Pathway enrichment was performed using Signaling Pathway Impact Analysis (SPIA) implemented via the iLINCS platform. Non-relevant pathways, such as those associated with viral responses (e.g., COVID-19), were excluded. Pathways related to epithelial biology were retained for interpretation.

### Deconvolution Analysis

sc-RNAseq dataset from^15^ was used to generate a signature matrix which was employed to derive cell proportions in SPARC IBD samples using CIBERSORTx (https://cibersortx.stanford.edu). Cell proportion chart was made using *ggplot2* in R version (4.3.0).

### Ordinal Logistic Regression Analysis

An ordinal logistic regression model (proportional odds model, polr function in the R package MASS) was used to assess the relationship between a transcriptomic signature score and macroscopic appearance in SPARC IBD samples. The signature score was derived as the mean z-scored expression of differentially expressed genes. Models were adjusted for clinical and demographic covariates (e.g., sex, race, smoking status), as well as an epithelial marker–based score to account for tissue composition. Odds ratios (OR) and 95% confidence intervals (CI) were calculated by exponentiating regression coefficients.

### Spline curve analysis

The relationship between the degree of inflammation in SPARC IBD data or DSS treatment in mouse model data and gene expression (expressed as the Z score of CPM) was visualized using a generalized additive model (GAM). we used the geom_smooth() function in the R package ggplot2 with the following parameters:

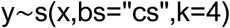

where y represents the Z score (derived from log2 normalized CPM values) and x represents the degree of inflammation. A cubic spline basis (bs = “cs”) was used with k=4 to control the smoothness of the fitted curve.

### DSS-mouse model

To model UC in mice, DSS or placebo was administered in the drinking water to induce acute colitis. C57BL/6 male mice aged 6–10 weeks were used. A solution of 2.5% DSS (w/v) was prepared using autoclaved water and provided to the mice for 6 days. Throughout the experiment, mice were monitored daily for body weight, stool consistency, and rectal bleeding. At the end-point, animals were euthanized, and colonic tissues were collected for histopathology assays. Adult mice were euthanized by inhalation of carbon dioxide (CO₂) for 6 minutes, delivered at a displacement rate of 20% of the cage volume per minute, in accordance with approved ethical protocols. No anaesthesia was used prior to euthanasia.

### PDOs establishment and culture

Human intestinal crypts were isolated from adult tissues using 25 μg/mL of liberase (Roche), embedded into Matrigel (Corning, Cat 356231) and grown into organoids following previous publications^75^. Organoids were routinely tested for mycoplasma contamination and resulted negative. The details of organoid donors used in each experiment are listed in Supplementary Table 4. Human intestinal organoids were cultured in IntestiCult™ Organoid Human Growth Medium or in-house media prepared following previous publications^75,76^ and were split once a week by mechanical dissociation and with TryPLE every two weeks.

### RNA extraction and sequencing of PDOs

RNA was collected from healthy individuals and IBD patient derived organoids embedded in Matrigel in 48-well plates. RNA was extracted using Qiagen RNeasy kit and RNA quality was assessed with the NanoDrop and the RIN score was checked. RNA samples were sequenced by Novogene, Cambridge, UK.

### Real-time PCR (qPCR)

250 ng of total RNA isolated from healthy and IBD-PDOs was converted to cDNA using Superscript II (Invitrogen). 20 µL of cDNA was diluted in 200 µL of nuclease free water. For analysing gene expression SYBR green (Applied biosystem) was used according to manufacturer’s protocol. All genes CT values were normalised with *GAPDH* (housekeeping gene) of each sample independently.

**Table 2.**
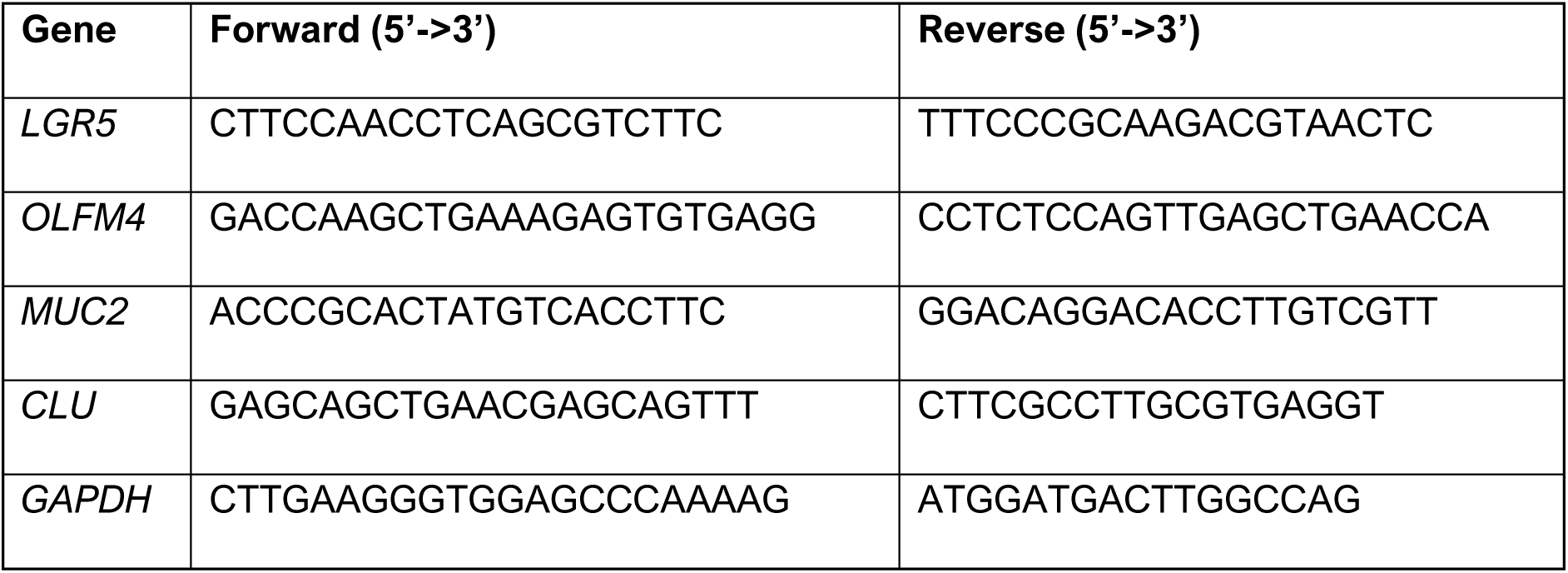
List of primers.

### Immunostaining

#### Paraffin processing from intestinal biopsies

Biopsy tissues were fixed in 4% PFA, placed into individual cassettes, and labelled accordingly. Tissue processing was performed using a Leica TP1020, enabling efficient handling of multiple samples. Tissues were exposed to a graded series of methanol solutions (50% to 100% concentration), with each step lasting 1 hour at RT under vacuum and agitation. Following dehydration, tissues were transferred to xylene for 1 hour at RT, again with vacuum and agitation, to remove residual alcohol and prepare the samples for paraffin infiltration. Embedding was carried out in molten paraffin using disposable molds (Embedding machine) while avoiding bubble formation. The embedded tissues were then cooled on the machine’s cooling surface until solidification, and the resulting paraffin blocks were subsequently stored at 4°C.

#### Tissue sectioning

A water bath was prepared with distilled water at 45°C for tissue sectioning. Meanwhile, the formalin-fixed paraffin-embedded (FFPE) blocks and forceps were chilled on ice. A microtome blade was installed onto the cutting bed with the correct orientation, and the FFPE block was secured in the block holder. Trimming was performed at 10-µm intervals until the desired depth was reached. Sections were then cut at 4 µm, forming a continuous ribbon that was gently transferred to the water bath using forceps. After briefly floating to separate the sections, slides (Superfrost) were used to mount the individual sections for subsequent Immunofluorescence (IF) staining.

#### Immunofluorescence staining

Paraffin-embedded tissue sections were placed on slides and dewaxed in xylene to remove residual paraffin. The sections were then rehydrated through sequential ethanol washes (100%, 90%, 80%, and 70%), each for 5 minutes, followed by distilled water. A 0.5 M sodium citrate buffer (pH 6.0) was freshly prepared, brought to a boil in a microwave, and the slides were submerged for 15 minutes before cooling to room temperature. Permeabilization was carried out using 0.5% Triton X-100 for 1 hour, and blocking solution was prepared by dissolving 0.1 g BSA in 10 ml of wash solution (0.1% Triton X-100 in PBS). Primary antibodies, diluted in blocking solution at the below described concentrations, were applied overnight at 4°C. Slides were washed three times, then incubated for 1 hour in the dark with secondary antibodies and DAPI (1:500 in blocking solution). After three additional washes using wash solution, coverslips were mounted using 100% glycerol. Imaging was performed using a Leica confocal SPE microscope.

#### Whole mount PDOs immunostaining

For immunofluorescence staining of organoids, we followed protocol from^77,78^ with slight modifications. Briefly, organoids were harvested in PBS for staining in 1.5 mL Protein Lowbind tube (Eppendorf) coated with 1% w/v BSA solution. Organoids were then fixed with 4% PFA for 1 hour at 4°C followed by centrifugation at 500 xg for 5 minutes. Permeabilization was performed using a solution of PBS with 0.5% Triton X-100 for 30 minutes at room temperature. Organoids were then blocked with a PBS solution containing 0.5% BSA for 30–60 minutes at 4°C. Primary antibody incubation was carried out overnight at 4°C on rotating platform using described concentration in table below. The following day, samples were washed three times with PBS and incubated with a secondary antibody mix containing DAPI for 1 hour at room temperature in the dark. After washing secondary antibody (dilution 1:500 of Alexa 488 anti-mouse or Alexa 568 anti-rabbit), organoids were plated in 96-well plate with black wall and clear bottom and imaged using Lecia Confocal SPE.

**Table 3.** List of antibodies.

| Antibodies and dyes | Company | Catalog | Dilution |
| --- | --- | --- | --- |
| S100A9 | Proteintech | 26992-1-AP | 1:200 |
| DAPI | Scientific Laboratory | D9542-1MG | 1:500 |
| OLFM4 | Cell Signalling | D6Y5A | 1:200 |
| β-catenin | Santacruz Biotech | sc-7963 | 1:500 |

## Data & code availability

The SPARC IBD data are available upon approved application to Crohn’s & Colitis Foundation IBD Plexus study (https://www.crohnscolitisfoundation.org/research/plexus). PDOs bulk-RNAseq data and the code done for this manuscript is deposited in Github Repository (https://github.com/brinda-bala/IBDstemcell.git).

## Declarations

### Ethics approval and consent to participate

Human tissue samples were collected in accordance with the Declaration of Helsinki and with approval from the East Midlands - Leicester Central Research Ethics Committee under reference number [17/EM/0126, protocol 17015]. Written informed consent was obtained from all participants prior to sample collection. *Approval date: 02/07/2020*. Study title: Modelling Intestinal iNflammation and predIctinG response to therapies Using human enTeroids

All animal experiments were performed in accordance with institutional guidelines and approved by the Cantonal Veterinary Office Vaud, Switzerland under license number [VD1052/VD1053]. Approval date: 1/11/2015. Study title: Production and analysis of mouse models of cancer stem cells

### Consent for publication

Not applicable

### Clinical trial number

Not applicable.

### Availability of data and material

The bulk RNA-seq data can be accessed here (GSE310845): https://www.ncbi.nlm.nih.gov/geo/query/acc.cgi?acc=GSE310845

### Authors’ contributions

P.O.M conceived, designed the study and wrote the manuscript with the help of B.B. P.O.M, B.B., and S.P. analysed and interpret data. B.B. carried out the modelling work, model-data comparison, RNAseq (single-cell and bulk) analyses, IBD Plexus analysis, experimental approaches and produced the figures together with S.P. W.D performed sc-RNAseq analyses by using Python. S.P performed mathematical tests, experimental approaches and transcriptomic (RNAseq and microarrays) analyses. G.M. compiled and helped with patients’ stratification, samples and clinical input. P.O.M, B.B, S.P., L.G., W.D., L.G., N.H., J.H., C.P., and G.M contributed to the discussions, interpretations and writing of the manuscript.

The authors declare that they have not use AI-generated work in this manuscript.

## Acknowledgements

We would like to express our gratitude to the patients who generously participated in this study and contributed their samples. We extend our sincere thanks to the nurses and clinical team from Queen’s Medical Centre (NIHR/NHS) Nottingham for their support and expertise in sample collection. Their commitment and professionalism were invaluable in facilitating this research. We would like to thank IBD Plexus team, US: the results published here are in part based on data from the Study of a Prospective Adult Research Cohort with IBD (SPARC IBD)

## Materials & Correspondence.

Correspondence and material requests should be addressed to P.O.M.

## References

1. GBD 2017 Inflammatory Bowel Disease Collaborators. The global, regional, and national burden of inflammatory bowel disease in 195 countries and territories, 1990-2017: a systematic analysis for the Global Burden of Disease Study 2017. Lancet Gastroenterol Hepatol 5, 17–30 (2020).

2. Banerjee, R., Raghunathan, N. & Pal, P. Managing inflammatory bowel disease: what to do when the best is unaffordable? The Lancet Gastroenterology & Hepatology 8, 396–398 (2023).

3. Lamb, C. A. et al. Inflammatory bowel disease has no borders: engaging patients as partners to deliver global, equitable and holistic health care. The Lancet 404, 414–417 (2024).

4. Gregorieff, A., Liu, Y., Inanlou, M. R., Khomchuk, Y. & Wrana, J. L. Yap-dependent reprogramming of Lgr5+ stem cells drives intestinal regeneration and cancer. Nature 526, 715–718 (2015).

5. Imajo, M., Ebisuya, M. & Nishida, E. Dual role of YAP and TAZ in renewal of the intestinal epithelium. Nat Cell Biol 17, 7–19 (2015).

6. Fevr, T., Robine, S., Louvard, D. & Huelsken, J. Wnt/beta-catenin is essential for intestinal homeostasis and maintenance of intestinal stem cells. Mol Cell Biol 27, 7551–7559 (2007).

7. Hollander, D. et al. Increased intestinal permeability in patients with Crohn’s disease and their relatives. A possible etiologic factor. Ann Intern Med 105, 883–885 (1986).

8. Turner, J. R. Intestinal mucosal barrier function in health and disease. Nat Rev Immunol 9, 799–809 (2009).

9. Dheer, R. et al. Intestinal Epithelial Toll-Like Receptor 4 Signaling Affects Epithelial Function and Colonic Microbiota and Promotes a Risk for Transmissible Colitis. Infect Immun 84, 798–810 (2016).

10. Barker, N. et al. Identification of stem cells in small intestine and colon by marker gene Lgr5. Nature 449, 1003–1007 (2007).

11. Capdevila, C. et al. Time-resolved fate mapping identifies the intestinal upper crypt zone as an origin of Lgr5+ crypt base columnar cells. Cell 187, 3039–3055.e14 (2024).

12. Malagola, E. et al. Isthmus progenitor cells contribute to homeostatic cellular turnover and support regeneration following intestinal injury. Cell 187, 3056–3071.e17 (2024).

13. Ito, G. et al. Lineage-Specific Expression of Bestrophin-2 and Bestrophin-4 in Human Intestinal Epithelial Cells. PLoS ONE 8, e79693 (2013).

14. Elmentaite, R. et al. Cells of the human intestinal tract mapped across space and time. Nature 597, 250–255 (2021).

15. Smillie, C. S. et al. Intra- and Inter-cellular Rewiring of the Human Colon during Ulcerative Colitis. Cell 178, 714–730.e22 (2019).

16. Parikh, K. et al. Colonic epithelial cell diversity in health and inflammatory bowel disease. Nature 567, 49–55 (2019).

17. Li, H. et al. Cross-species single-cell transcriptomic analysis reveals divergence of cell composition and functions in mammalian ileum epithelium. Cell Regen 11, 19 (2022).

18. Širvinskas, D. et al. Single-cell atlas of the aging mouse colon. iScience 25, 104202 (2022).

19. Tsang, D. K. L. et al. A single cell survey of the microbial impacts on the mouse small intestinal epithelium. Gut Microbes 14, 2108281 (2022).

20. Schumacher, M. A. et al. Deep Crypt Secretory Cell Differentiation in the Colonic Epithelium Is Regulated by Sprouty2 and Interleukin 13. Cellular and Molecular Gastroenterology and Hepatology 15, 971–984 (2023).

21. Raffals, L. E. et al. The Development and Initial Findings of A Study of a Prospective Adult Research Cohort with Inflammatory Bowel Disease (SPARC IBD). Inflamm Bowel Dis 28, 192–199 (2022).

22. Gassler, N. et al. Inflammatory bowel disease is associated with changes of enterocytic junctions. American Journal of Physiology-Gastrointestinal and Liver Physiology 281, G216–G228 (2001).

23. Van Der Flier, L. G., Haegebarth, A., Stange, D. E., Van De Wetering, M. & Clevers, H. OLFM4 Is a Robust Marker for Stem Cells in Human Intestine and Marks a Subset of Colorectal Cancer Cells. Gastroenterology 137, 15–17 (2009).

24. Ayyaz, A. et al. Single-cell transcriptomes of the regenerating intestine reveal a revival stem cell. Nature 569, 121–125 (2019).

25. Fang, K. et al. Temporal genomewide expression profiling of DSS colitis reveals novel inflammatory and angiogenesis genes similar to ulcerative colitis. Physiological Genomics 43, 43–56 (2011).

26. Hong, D. et al. Integrative analysis of single-cell RNA-seq and gut microbiome metabarcoding data elucidates macrophage dysfunction in mice with DSS-induced ulcerative colitis. Commun Biol 7, 731 (2024).

27. Korndörfer, I. P., Brueckner, F. & Skerra, A. The crystal structure of the human (S100A8/S100A9)2 heterotetramer, calprotectin, illustrates how conformational changes of interacting alpha-helices can determine specific association of two EF-hand proteins. J Mol Biol 370, 887–898 (2007).

28. Nanki, K. et al. Somatic inflammatory gene mutations in human ulcerative colitis epithelium. Nature 577, 254–259 (2020).

29. Sato, T. et al. Long-term expansion of epithelial organoids from human colon, adenoma, adenocarcinoma, and Barrett’s epithelium. Gastroenterology 141, 1762–1772 (2011).

30. Ootani, A. et al. Sustained in vitro intestinal epithelial culture within a Wnt-dependent stem cell niche. Nat Med 15, 701–706 (2009).

31. Dotti, I. et al. Alterations in the epithelial stem cell compartment could contribute to permanent changes in the mucosa of patients with ulcerative colitis. Gut 66, 2069–2079 (2017).

32. Burclaff, J. et al. A Proximal-to-Distal Survey of Healthy Adult Human Small Intestine and Colon Epithelium by Single-Cell Transcriptomics. Cellular and Molecular Gastroenterology and Hepatology 13, 1554–1589 (2022).

33. Patnaude, L. et al. Mechanisms and regulation of IL-22-mediated intestinal epithelial homeostasis and repair. Life Sci 271, 119195 (2021).

34. Cineus, R. et al. The IL-22–oncostatin M axis promotes intestinal inflammation and tumorigenesis. Nat Immunol 26, 837–853 (2025).

35. He, G.-W. et al. Optimized human intestinal organoid model reveals interleukin-22-dependency of paneth cell formation. Cell Stem Cell 29, 1333–1345.e6 (2022).

36. Lindemans, C. A. et al. Interleukin-22 promotes intestinal-stem-cell-mediated epithelial regeneration. Nature 528, 560–564 (2015).

37. Kong, L. et al. The landscape of immune dysregulation in Crohn’s disease revealed through single-cell transcriptomic profiling in the ileum and colon. Immunity 56, 444–458.e5 (2023).

38. Aran, D. et al. Reference-based analysis of lung single-cell sequencing reveals a transitional profibrotic macrophage. Nat Immunol 20, 163–172 (2019).

39. Vong, L. et al. Up-regulation of Annexin-A1 and lipoxin A(4) in individuals with ulcerative colitis may promote mucosal homeostasis. PLoS One 7, e39244 (2012).

40. Nanakin, A. et al. Expression of the REG IV gene in ulcerative colitis. Laboratory Investigation 87, 304–314 (2007).

41. Kapsoritakis, A. N. et al. Imbalance of tissue inhibitors of metalloproteinases (TIMP) – 1 and – 4 serum levels, in patients with inflammatory bowel disease. BMC Gastroenterol 8, 55 (2008).

42. Hensel, K. O. et al. Differential expression of mucosal trefoil factors and mucins in pediatric inflammatory bowel diseases. Sci Rep 4, 7343 (2014).

43. Okayasu, I. et al. A novel method in the induction of reliable experimental acute and chronic ulcerative colitis in mice. Gastroenterology 98, 694–702 (1990).

44. Ungaro, R., Mehandru, S., Allen, P. B., Peyrin-Biroulet, L. & Colombel, J.-F. Ulcerative colitis. The Lancet 389, 1756–1770 (2017).

45. Torres, J. et al. Serum Biomarkers Identify Patients Who Will Develop Inflammatory Bowel Diseases Up to 5 Years Before Diagnosis. Gastroenterology 159, 96–104 (2020).

46. Honap, S. et al. Prevention and interception trials in inflammatory bowel disease: an international taskforce assessment on clinical trial design. Lancet Gastroenterol Hepatol S2468-1253(24)00439–4 (2025) doi:10.1016/S2468-1253(24)00439-4.

47. Gomollón, F. et al. 3rd European Evidence-based Consensus on the Diagnosis and Management of Crohn’s Disease 2016: Part 1: Diagnosis and Medical Management. ECCOJC 11, 3–25 (2017).

48. Sands, B. E. & Ooi, C. J. A survey of methodological variation in the Crohn’s disease activity index. Inflamm Bowel Dis 11, 133–138 (2005).

49. Sanman, L. E. et al. Transit-Amplifying Cells Coordinate Changes in Intestinal Epithelial Cell-Type Composition. Dev Cell 56, 356–365.e9 (2021).

50. Parigi, S. M. et al. The spatial transcriptomic landscape of the healing mouse intestine following damage. Nat Commun 13, 828 (2022).

51. Heuberger, J. et al. Shp2/MAPK signaling controls goblet/paneth cell fate decisions in the intestine. Proc. Natl. Acad. Sci. U.S.A. 111, 3472–3477 (2014).

52. Kabiri, Z. et al. Wnt signaling suppresses MAPK-driven proliferation of intestinal stem cells. Journal of Clinical Investigation 128, 3806–3812 (2018).

53. Grivennikov, S. et al. IL-6 and Stat3 are required for survival of intestinal epithelial cells and development of colitis-associated cancer. Cancer Cell 15, 103–113 (2009).

54. Pickert, G. et al. STAT3 links IL-22 signaling in intestinal epithelial cells to mucosal wound healing. J Exp Med 206, 1465–1472 (2009).

55. Herrera, S. C. & Bach, E. A. JAK/STAT signaling in stem cells and regeneration: from Drosophila to vertebrates. Development 146, dev167643 (2019).

56. Biton, M. et al. T Helper Cell Cytokines Modulate Intestinal Stem Cell Renewal and Differentiation. Cell 175, 1307–1320.e22 (2018).

57. Van Unen, V. et al. Identification of a Disease-Associated Network of Intestinal Immune Cells in Treatment-Naive Inflammatory Bowel Disease. Front. Immunol. 13, 893803 (2022).

58. Oliver, A. J. et al. Single-cell integration reveals metaplasia in inflammatory gut diseases. Nature 635, 699–707 (2024).

59. Planell, N. et al. Transcriptional analysis of the intestinal mucosa of patients with ulcerative colitis in remission reveals lasting epithelial cell alterations. Gut 62, 967–976 (2013).

60. Friedrich, M. et al. IL-1-driven stromal-neutrophil interactions define a subset of patients with inflammatory bowel disease that does not respond to therapies. Nat Med 27, 1970–1981 (2021).

61. Li, J. et al. Identification and multimodal characterization of a specialized epithelial cell type associated with Crohn’s disease. Nat Commun 15, 7204 (2024).

62. Arato, A., Savilahti, E. & Paszti, I. Crypt hyperplasia related to increased lymphocyte activation in the rectal mucosa of children with ulcerative colitis. Z Gastroenterol 32, 483–487 (1994).

63. Buczacki, S. J. A. et al. Intestinal label-retaining cells are secretory precursors expressing Lgr5. Nature 495, 65–69 (2013).

64. Nowell, C. S. et al. Chronic inflammation imposes aberrant cell fate in regenerating epithelia through mechanotransduction. Nat Cell Biol 18, 168–180 (2016).

65. Neurath, M. F., Pettersson, S., Meyer zum Büschenfelde, K. H. & Strober, W. Local administration of antisense phosphorothioate oligonucleotides to the p65 subunit of NF-kappa B abrogates established experimental colitis in mice. Nat Med 2, 998–1004 (1996).

66. Rogler, G. et al. Nuclear factor kappaB is activated in macrophages and epithelial cells of inflamed intestinal mucosa. Gastroenterology 115, 357–369 (1998).

67. Schwitalla, S. et al. Intestinal Tumorigenesis Initiated by Dedifferentiation and Acquisition of Stem-Cell-like Properties. Cell 152, 25–38 (2013).

68. Myant, K. B. et al. ROS Production and NF-κB Activation Triggered by RAC1 Facilitate WNT-Driven Intestinal Stem Cell Proliferation and Colorectal Cancer Initiation. Cell Stem Cell 12, 761–773 (2013).

69. Jones, L. G. et al. NF-κB2 signalling in enteroids modulates enterocyte responses to secreted factors from bone marrow-derived dendritic cells. Cell Death Dis 10, 896 (2019).

70. Burkitt, M. D. et al. NF-κB1, NF-κB2 and c-Rel differentially regulate susceptibility to colitis-associated adenoma development in C57BL/6 mice. J Pathol 236, 326–336 (2015).

71. Guo, Q. et al. NF-κB in biology and targeted therapy: new insights and translational implications. Sig Transduct Target Ther 9, 53 (2024).

72. Yunshun Chen <Yuchen@Wehi. Edu.Au>, A. L. E. A. edgeR. Bioconductor 10.18129/B9.BIOC.EDGER (2017).

73. Gordon Smyth [Cre, A. limma. Bioconductor 10.18129/B9.BIOC.LIMMA (2017).

74. Hao, Y. et al. Dictionary learning for integrative, multimodal and scalable single-cell analysis. Nat Biotechnol 42, 293–304 (2024).

75. Ordóñez-Morán, P., Dafflon, C., Imajo, M., Nishida, E. & Huelsken, J. HOXA5 Counteracts Stem Cell Traits by Inhibiting Wnt Signaling in Colorectal Cancer. Cancer Cell 28, 815–829 (2015).

76. Hacker, D. L. & Ordóñez-Morán, P. Large-Scale Production of Recombinant Noggin and R-Spondin1 Proteins Required for the Maintenance of Stem Cells in Intestinal Organoid Cultures. Methods Mol Biol 2171, 171–184 (2020).

77. Norkin, M., Ordóñez-Morán, P. & Huelsken, J. High-content, targeted RNA-seq screening in organoids for drug discovery in colorectal cancer. Cell Rep 35, 109026 (2021).

78. Pleguezuelos-Manzano, C. et al. Establishment and Culture of Human Intestinal Organoids Derived from Adult Stem Cells. Curr Protoc Immunol 130, e106 (2020).

