## Supplementary material for "Intestinal Stem and Progenitor Cells Exhibit Distinct Adaptive Responses to Inflammatory Stress in IBD": Supp

Supplementary Figure 1

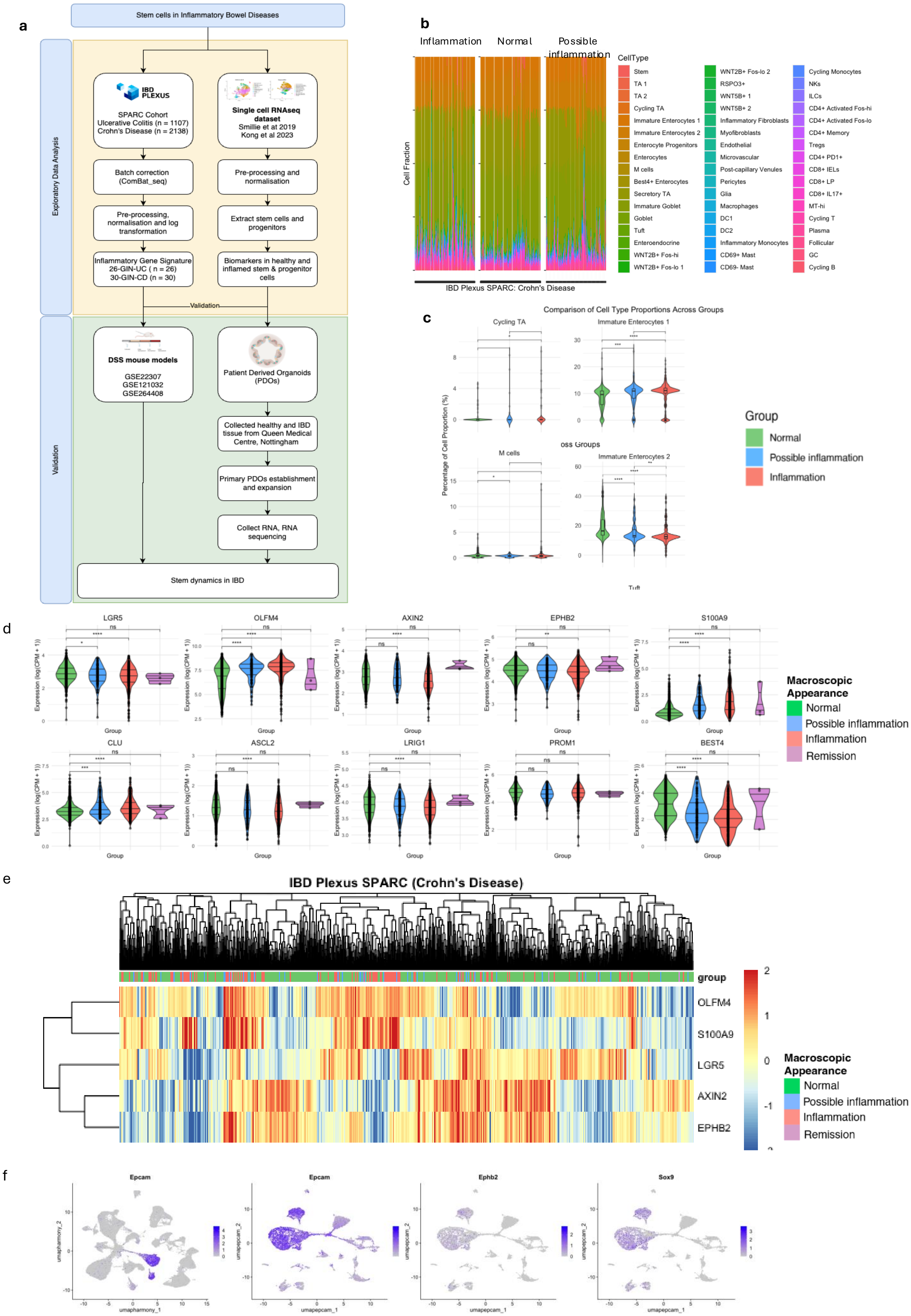

**Figure S1.** Epithelial cell heterogeneity profile based on inflammation severity.
a) Schematic overview of the bioinformatics and experimental study workflow.
b) Bar plots showing the proportion of epithelial cell types across SPARC IBD-CD samples, stratified by endoscopic macroscopic assessment.
c) Violin plots histograms showing the significant changes in cell type proportions according to macroscopic appearance.
d) Violin plots showing comparative gene expression markers according to macroscopic appearance.
e) Heatmap showing unsupervised clustering of gene expression markers according to macroscopic appearance.
f) EPCAM+ epithelial cells were extracted from the total cell population of sc-RNAseq DSS-treated mice with normal, acute and chronic colitis conditions (GSE264408) dataset.

Supplementary Figure 2

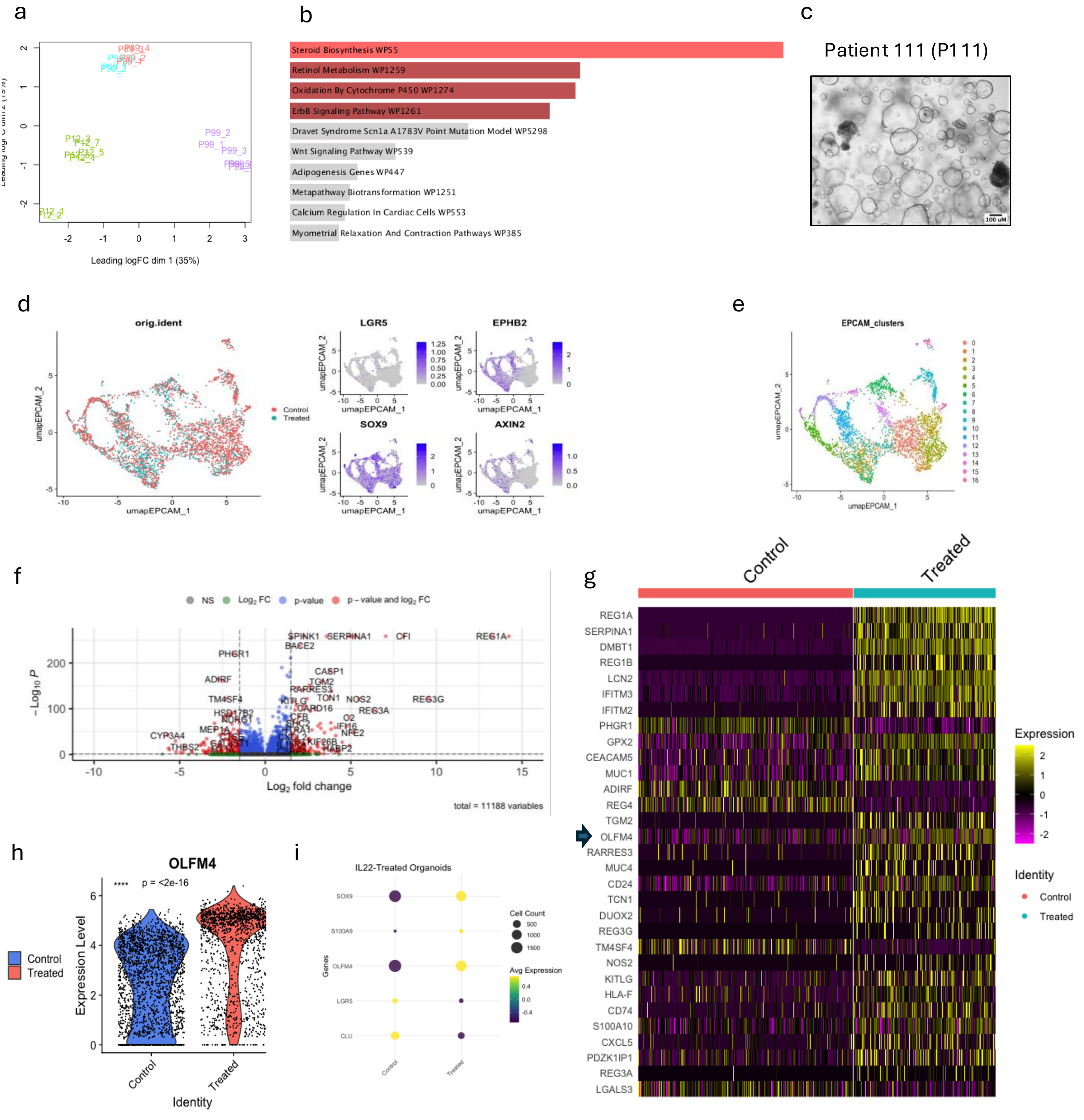


**Figure S2.** Characterization of healthy and IBD epithelial cells using a PDO model.

a) PCA plot showing the sample distribution of bulk-RNAseq samples of PDOs: hP12 in green (early and late passages), IBD-P89 in red, IBD-P90 in blue, and IBD-P99 in purple colour (early and late passages).

b) Enrichment analysis of downregulated IBD-PDOs vs hPDOs DEGs

c) Representative image of additional IBD-PDO included to perform qPCR analysis

d) UMAPs showing the sc-RNAseq EPCAM$^+$ cells of IL-22 treated and control hPDOs (left), *LGR5*, *EPHB2*, *SOX9* and *AXIN2* gene expression (middle), and e) clusters from 1 to 16 (right) (GSE189423).

f) Volcano plot showing the DEGs in EPCAM$^+$ IL-22 treated and control hPDOs as red dots (adjusted $p < 0.05$, log2 fold change cutoff at 1.5) and non-significant genes as grey dots (GSE189423).

g) Heatmap showing expression levels of DEGs in EPCAM$^+$ IL-22 treated and control hPDOs (GSE189423).

h) Violin plot showing gene expression levels of *OLFM4* in IL-22 treated and control hPDOs (GSE189423).

i) Dot plot showing gene expression markers according to sc-RNAseq data of DSS-treated mice categorised by normal, acute and chronic colitis (GSE264408).

Supplementary Figure 3

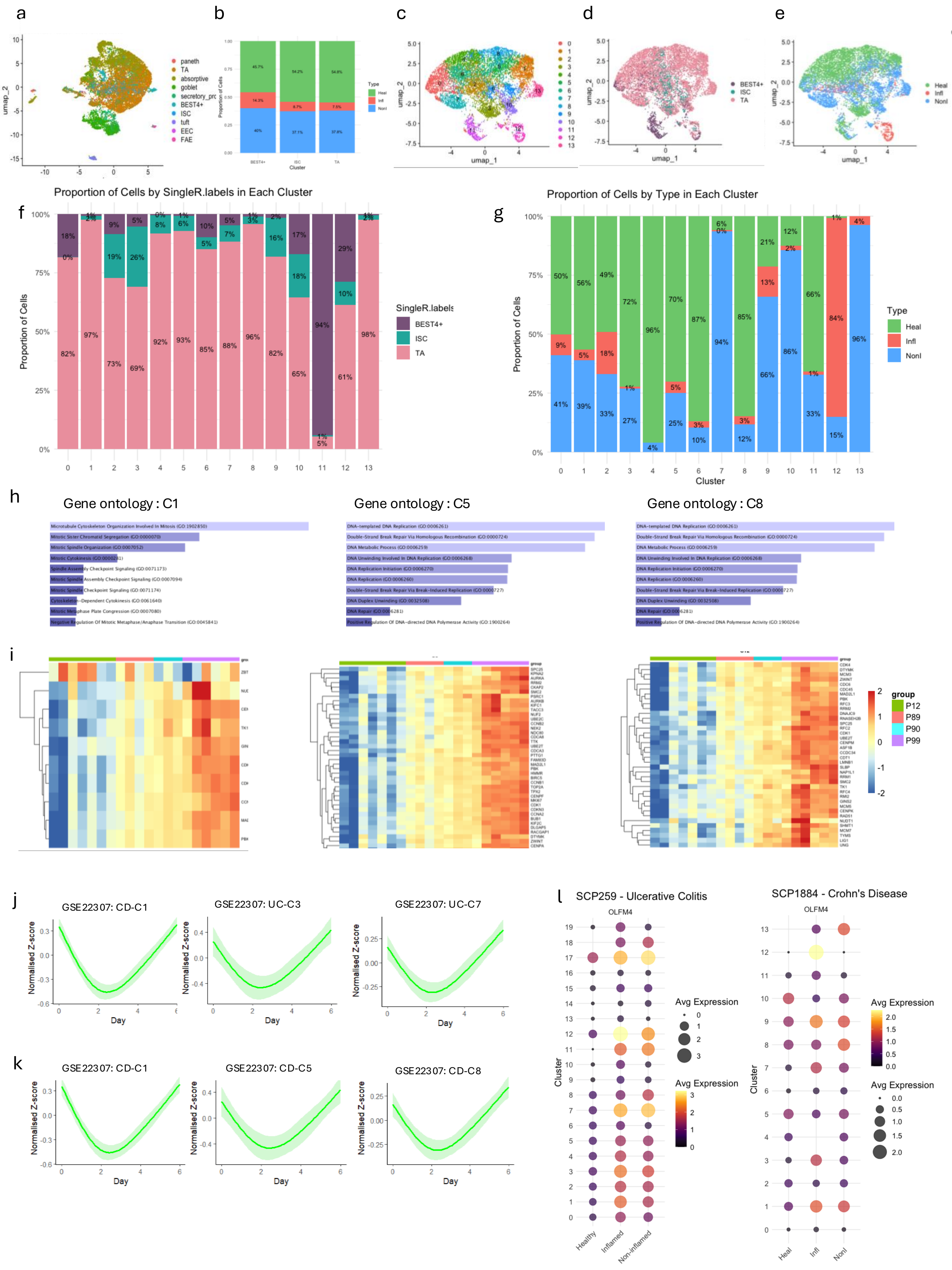

**Figure S3.** SC/TA and BEST4⁺ pools in healthy and CD non-inflamed and inflamed tissue.
a) UMAP of the cell type annotations predicted in CD dataset including SC, TA, and BEST4⁺ pools (Burclaff et al 2022).
b) Bar plot showing the quantification of SC/TA, and BEST4⁺ cells derived from inflamed, non-inflamed and normal tissue (SCP1884).
c) UMAP plot showing 14 distinct transcriptional clusters (CD-C0 to CD-C13) identified in the CD dataset, highlighting cellular heterogeneity within the SC/TA, and BEST4⁺ populations.
UMAP plot by d) cell type and e) health status.
f) Bar plot of proportional distribution of clusters based on health status.
g) Bar plot of proportional distribution of clusters based on cell type.
h) Bar plots showing the enriched pathways from Gene Ontology (GO-Biological Processes 2023) of CD-C1, -C5 and –C8 markers.
i) Heatmaps of UC-C1, -C3 and -C7 expression markers in healthy and IBD-PDOs.
j) Spline curves showing the dynamic shift in expression levels of UC-C1, -C3 and -C7 markers in DSS-treated mice (GSE22307).
k) Spline curves showing the dynamic shift in expression levels of CD-C1, -C5 and -C8 markers in DSS-treated mice (GSE22307).
l) Bubble plots showing *OLFM4* expression in the different UC and CD clusters.
.

Supplementary Figure 4

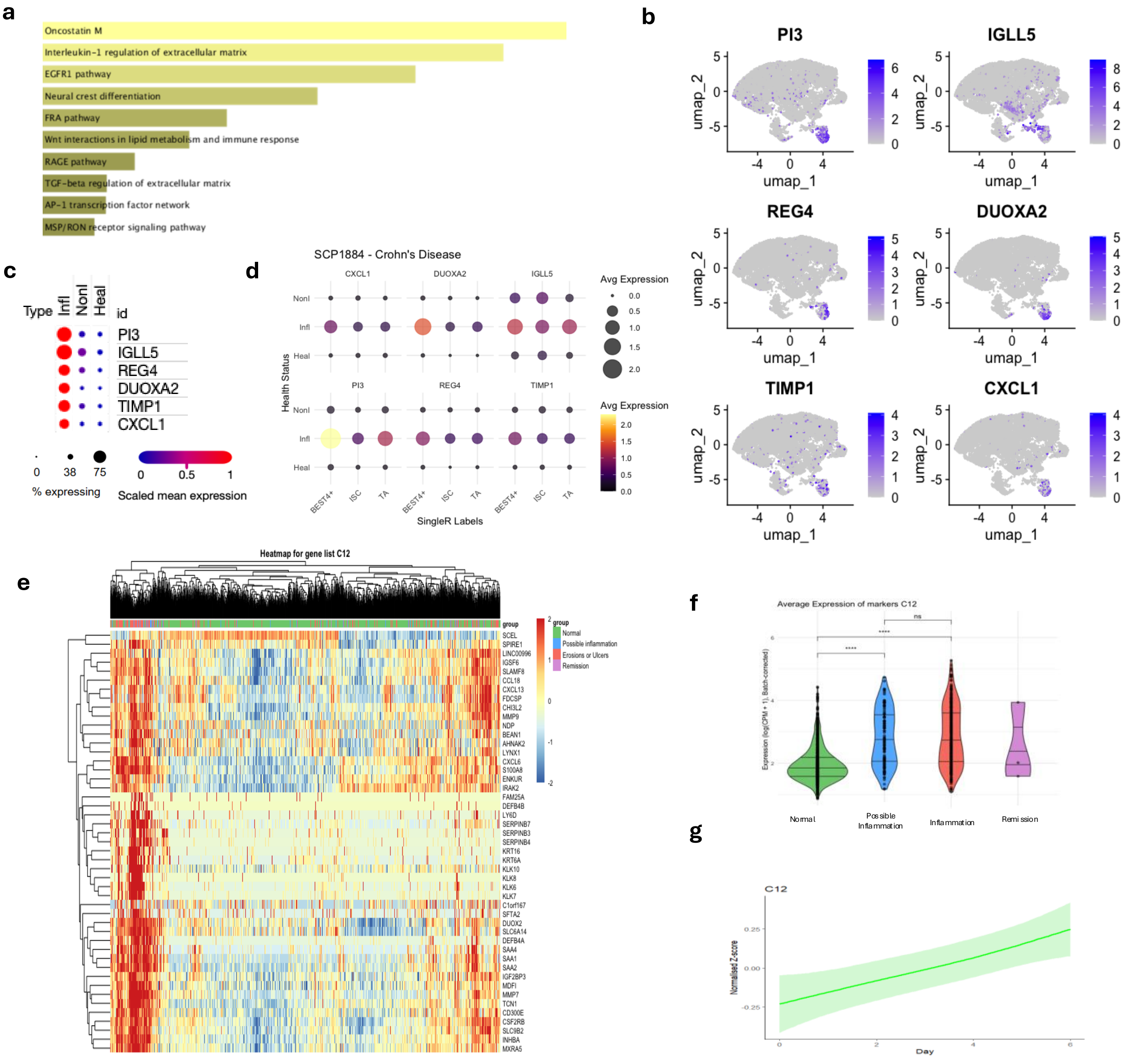

**Figure S4.** Immune-enriched gene clusters in CD SC/TA and BEST4+ pools during inflammation.
a) Bar plots showing the Gene Ontology (Biological Processes) enriched pathways of CD-C12 50 top genes.
b) Feature plot showing *PI3*, *IGLL5*, *REG4*, *DUOXA2*, *TIMP1* and *CXCL1* expression in sc-RNAseq SC/TA and BEST4+ clusters (SCP1884).
c) Dot plot showing *PI3*, *IGLL5*, *REG4*, *DUOXA2*, *TIMP1* and *CXCL1* expression in epithelial cells according to tissue source categorised by non-inflamed, inflamed and healthy groups (SCP1884).
d) Dot plot showing *PI3*, *IGLL5*, *REG4*, *DUOXA2*, *TIMP1* and *CXCL1* expression in BEST4+, SC and TA cell types according to tissue source categorised by non-inflamed, inflamed and healthy groups (SCP1884).
e) Heatmap showing the unsupervised clustering of CD-C12 gene expression markers in SPARC IBD-CD stratified by macroscopic appearance (normal, possible inflammation and inflammation).
f) Violin plots histograms of CD-C12 genes expression categorised by SPARC IBD-CD macroscopic appearance (normal, possible inflammation and inflammation).
g) Average expression spline curves of the CD-C12 50 top genes of DSS-treated mice dataset (GSE22307).

Supplementary Figure 5

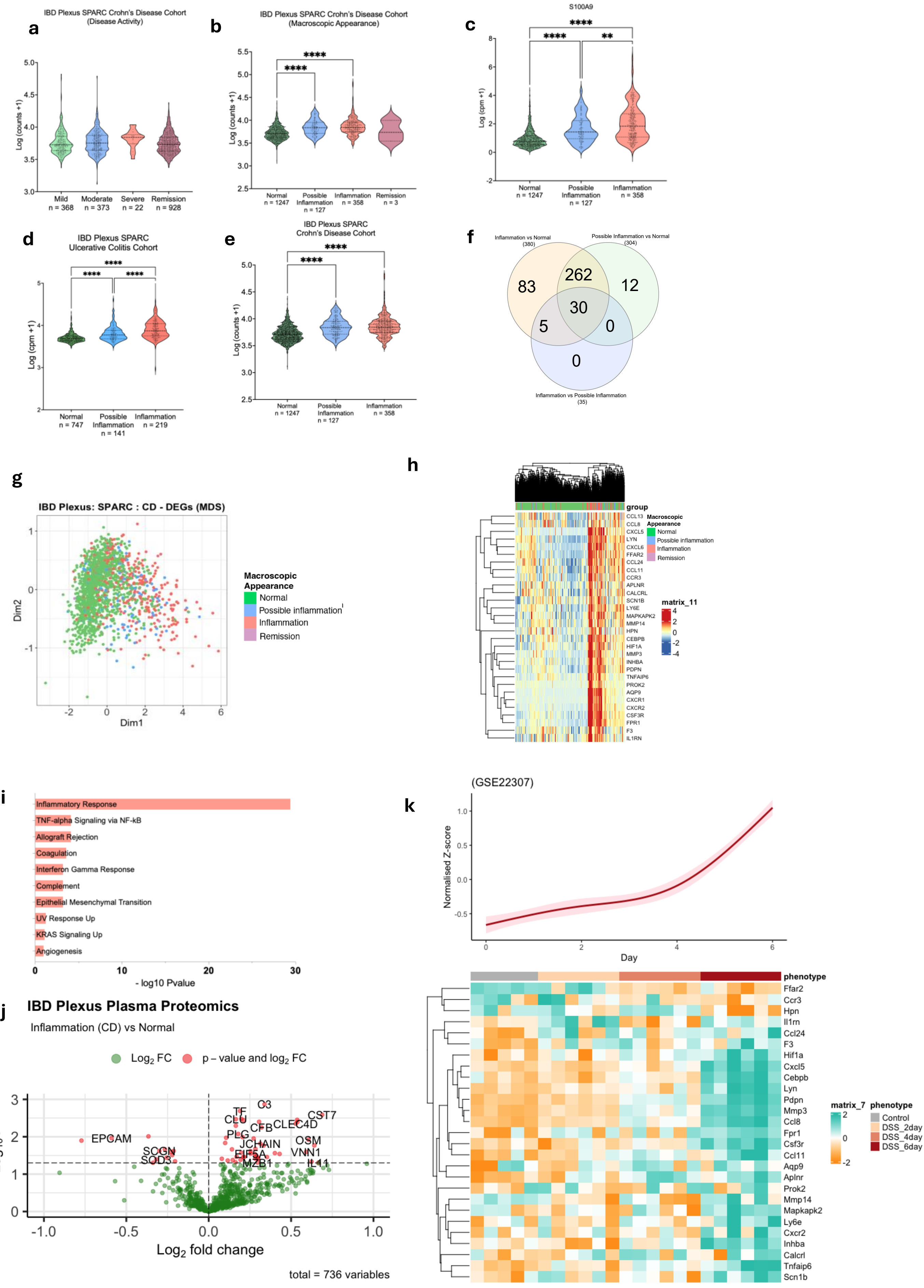

**Figure S5.** A novel inflammation-linked gene signature associated with inflammation severity in IBD.
Violin plot histograms represent the comparative average gene expression of curated inflammatory genes (n = 476) in SPARC IBD-CD samples categorised by: Bar plots showing the proportion of cells in each cell type per SPARC IBD-CD endoscopic assessment category.
a) disease activity, b) macroscopic appearance and c) *S100A9* in normal (n = 1,247), possible inflammation (n = 127), inflammation (n = 358) and remission (n = 3) groups.
Violin plot histogram showing the comparative gene expression of a public available inflammatory gene signature (Smilie et al, 2019) in d) UC and e) CD.
f) Venn diagram showing the overlap of significantly DEGs to identify the novel gene signature (30-GIN-CD).
g) Multidimensionality Scaling (MDS) sample distribution according to gene expression profile within the SPARC IBD-CD cohort in normal, possible inflammation and inflammation groups.
h) Heatmap represents unsupervised clustering of the 30-GIN-CD genes within the SPARC IBD-CD cohort.
i) Significantly enriched pathways obtained by Enrichr (MSigDB) of the 30-GIN-CD genes.
j) Volcano plot illustrate significantly differentially expressed proteins (DEPs) in the inflammation vs normal groups as red dots (adjusted $p < 0.05$, log2 fold changes) and non-significant proteins as green.
k) Heatmaps and average expression spline curves (above) depicting the dynamic expression shifts of the 30-GIN-CD in DSS-treated mice (GSE22307).
** $p < 0.01$, **** $p < 0.0001$

### Supplementary Figure 6

**a** Positive correlation of SPARC IBD UC individual gene expression with UC-C18 gene markers

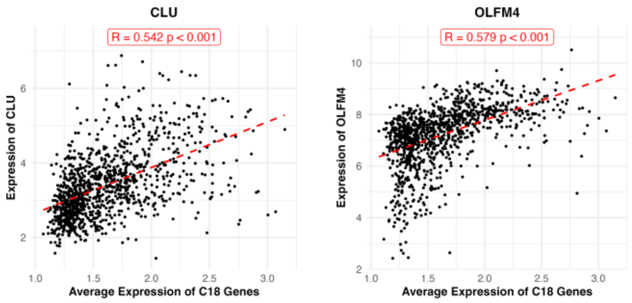


**b** Weak negative or no correlation of SPARC IBD UC individual gene expression with UC-C18 gene markers

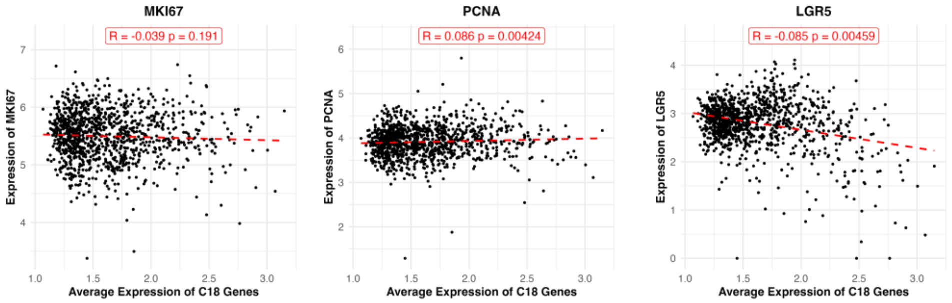


**c** Negative corelation of SPARC IBD UC individual gene expression with UC-C18 gene markers

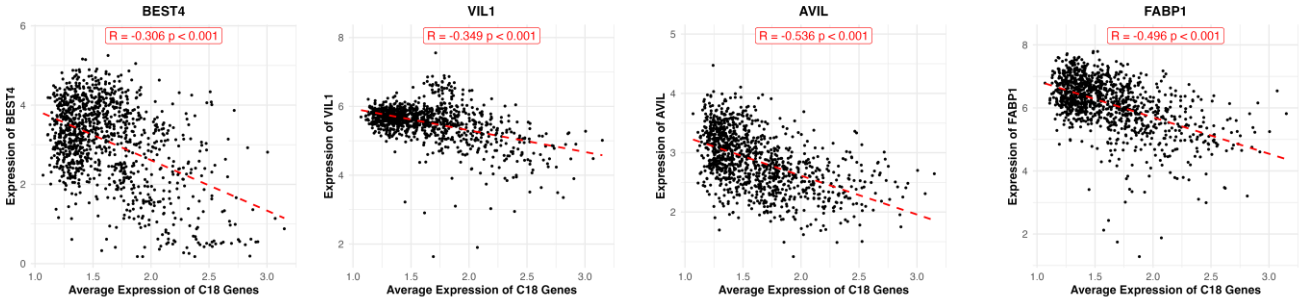


**Figure S6.**
Scatter Plots showing the:
a) positive correlation of *CLU* and *OLFM4*
b) weak negative or no correlation of *MKI67*, *PCNA* and *LGR5*
c) negative correlation of *BEST4, VIL1*, *AVIL*, and *FABP1*.
with UC-C18 gene markers in SPARC IBD -UC, respectively

### Supplementary Figure 7

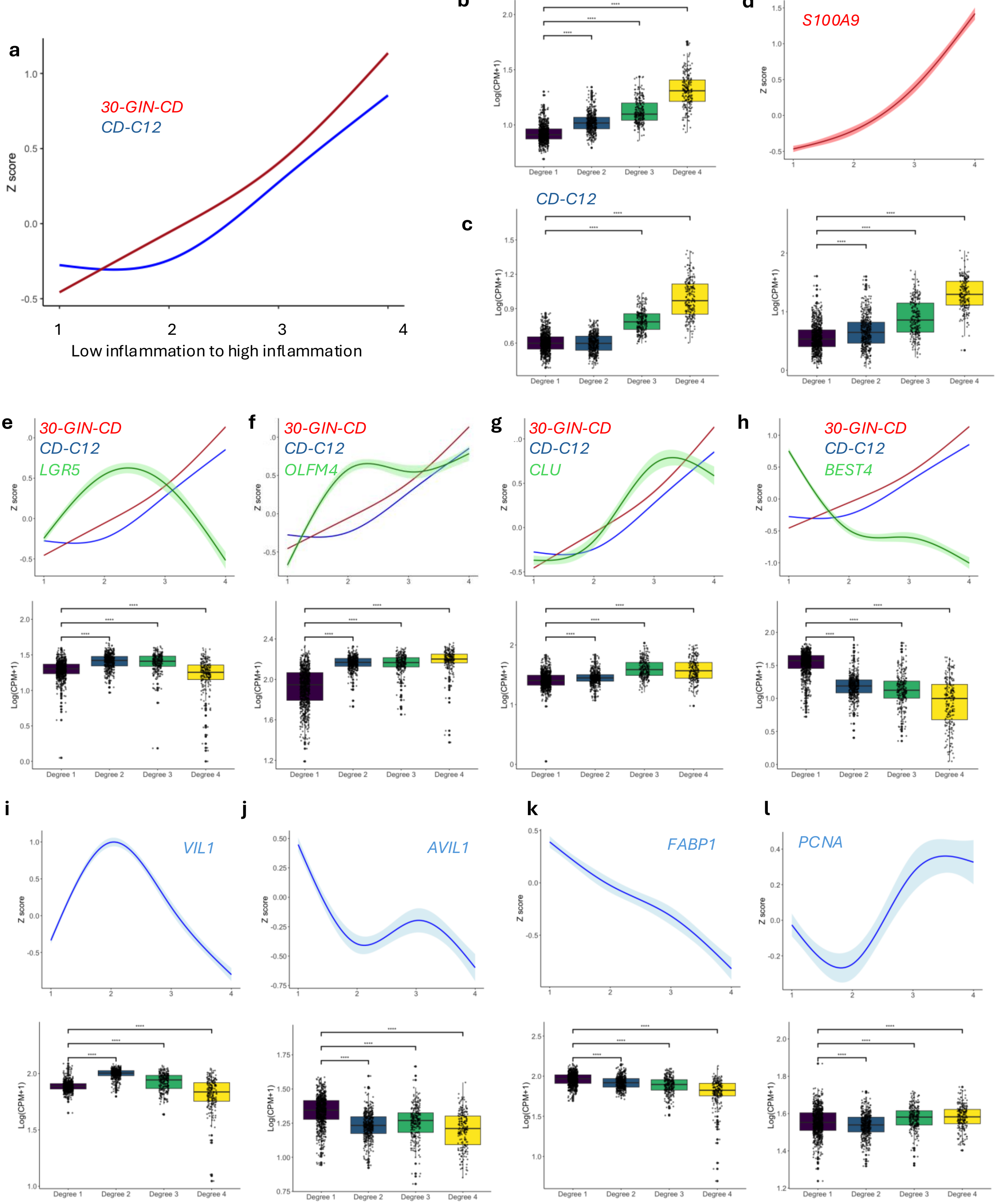


**Figure S7.** Adaptive dynamics of epithelial cells in CD tissue depending on the inflammation degree.
Spline curve (a) and box plot of 30-GIN-CD (b) and CD-C12 (c) gene signatures in low to high inflammation degree.
d) Spline curve (top) and box plot (bottom) of *S100A9* expression in low to high inflammation degree.
Spline curves (top) and box plots (bottom) of CD-C12, 30-GIN-CD and individual gene expression of e) *LGR5*, f) *OLFM4*, g) *CLU*, h) *BEST4* in low to high inflammation degree.
Spline curves (top) and box plots (bottom) of I) *VIL1*, j) *AVIL1*, k) *FABP1*, and l) *PCNA* in low to high inflammation degree.

Supplementary Figure 7 (cont.)

**m** Positive correlation of SPARC IBD CD data with CD-C12 markers

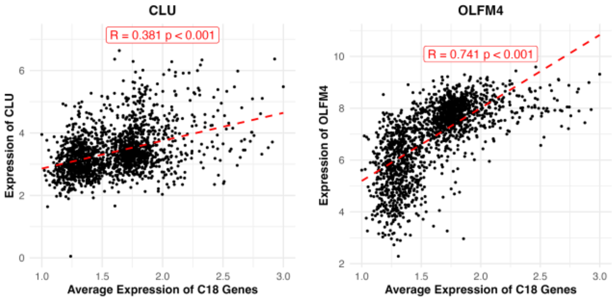


**n** Weak correlation or no correlation of SPARC IBD CD data with CD-C12 markers

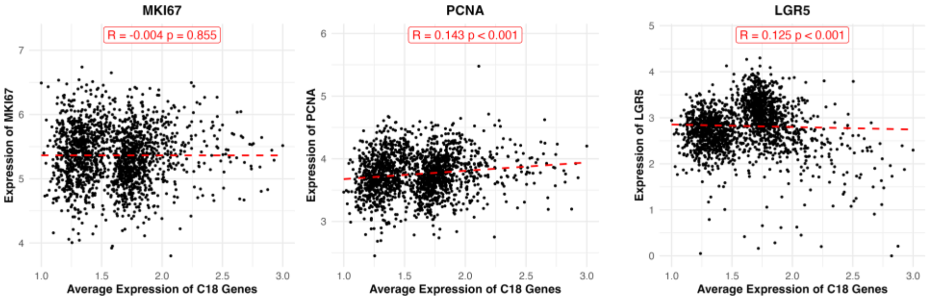


**o** Negative correlation of SPARC IBD CD data with CD-C12 markers

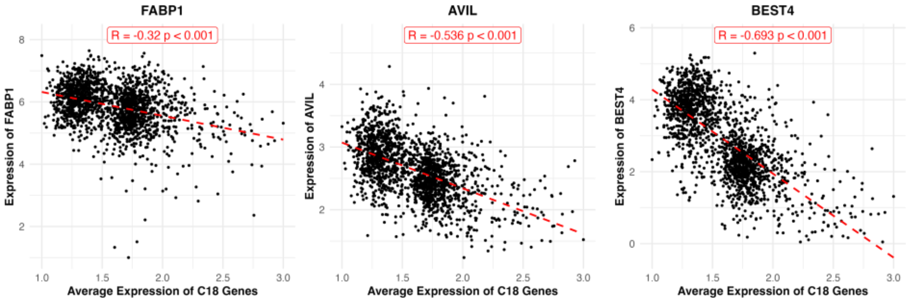


**Figure S7 cont.**
Scatter Plots showing the:
m) Positive correlation of *CLU* and *OLFM4*
n) Weak negative or no correlation of *MKI67*, *PCNA* and *LGR5*
o) Negative correlation of *FABP1, AVIL*, and *BEST4*
with CD-C12 markers in SPARC IBD CD cohort, respectively

Supplementary Figure 7 (cont.)

**p**

IBD Plexus SPARC –
Ulcerative Colitis Cohort

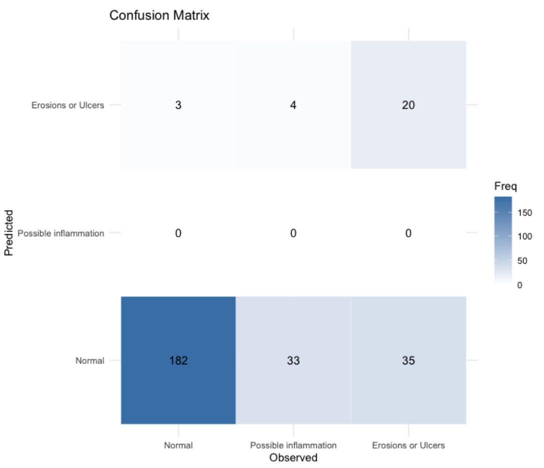


**q**

IBD Plexus SPARC –
Crohn's Disease Cohort

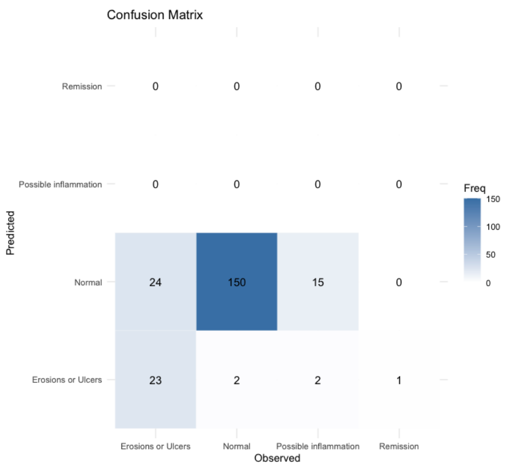


p) Model performance was evaluated using a patient-level hold-out set (75% training, 25% testing), and predictive accuracy was summarized with confusion matrix done for UC data (above) and
q) CD data (below).

### Supplementary Figure 8

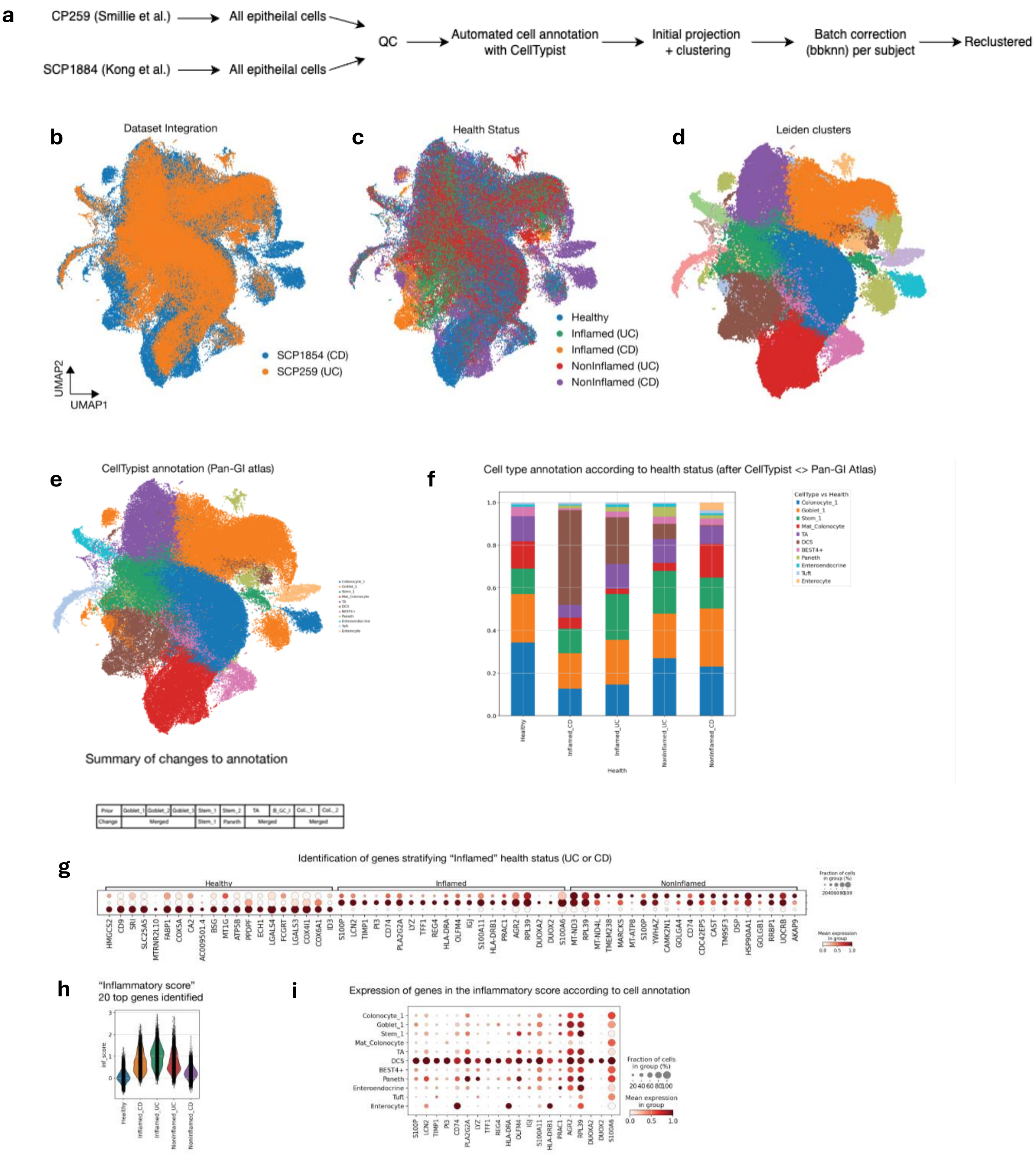


**Figure S8.** Single-cell RNA-seq data integration analysis.
a) Schematic overview of the integrated single-cell RNA-seq analysis pipeline.
b) UMAP showing the distribution of all epithelial cells across datasets.
c) UMAP visualisation coloured by health status (healthy, non-inflamed IBD, inflamed UC/CD).
d) UMAP showing Leiden clustering results across the integrated dataset.
e) Summary table displaying automated Pan-GI cell type labels and manual refinements to cell identity annotations.
f) Stacked bar plot illustrating cell type distribution by health status, highlighting the loss of colonocytes and expansion of deep crypt secretory (DCS) cells during inflammation.
g) Bubble plot showing the expression of inflammation-associated genes across health status categories.
h) Violin plot showing inflammatory gene signature scores (top 20 genes) elevated in inflamed UC and CD samples compared to non-inflamed IBD and healthy tissues.
i) Bubble plot illustrating the expression of the top 20 inflammation-associated genes across annotated epithelial cell types.

#### Supplementary Figure 8 (cont.)

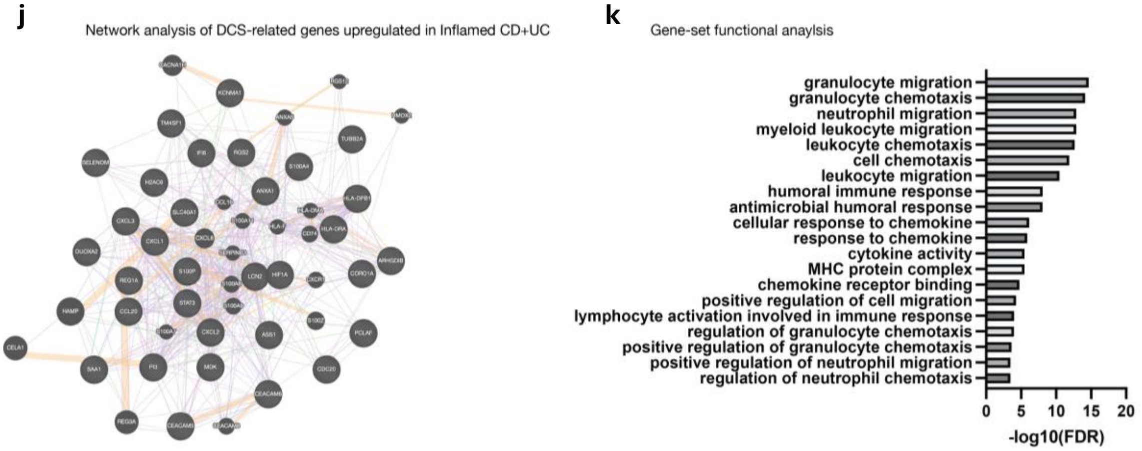


**Figure S8 (cont.)**

j) Network analysis of dendritic cell signature (DCS)-related genes significantly upregulated in inflamed Crohn's disease (CD) and ulcerative colitis (UC) tissue. The network illustrates key interactions among these genes, highlighting potential hubs involved in inflammatory responses.

k) Gene-set functional enrichment analysis of the network-derived gene clusters showing significant enrichment of inflammation-associated pathways and gene ontology terms, supporting the involvement of these genes in immune and inflammatory processes in CD and UC.
